# Phased ERK-responsiveness and developmental robustness regulate teleost skin morphogenesis

**DOI:** 10.1101/2024.05.13.593750

**Authors:** Nitya Ramkumar, Christian Richardson, Makinnon O’Brien, Faraz Ahmed Butt, Jieun Park, Anna T Chao, Michel Bagnat, Kenneth Poss, Stefano Di Talia

## Abstract

Elongation of the vertebrate embryonic axis necessitates rapid expansion of the epidermis to accommodate the growth of underlying tissues. Here, we generated a toolkit to visualize and quantify signaling in entire cell populations of periderm, the outermost layer of the epidermis, in live developing zebrafish. We find that oriented cell divisions facilitate growth of the early periderm during axial elongation rather than cell addition from the basal layer. Activity levels of ERK, a downstream effector of MAPK pathway, gauged by a live biosensor, predicts cell cycle entry, and optogenetic ERK activation controls proliferation dynamics. As development proceeds, rates of peridermal cell proliferation decrease, ERK activity becomes more pulsatile and functionally transitions to promote hypertrophic cell growth. Targeted genetic blockade of cell division generates animals with oversized periderm cells, yet, unexpectedly, development to adulthood is not impaired. Our findings reveal stage-dependent differential responsiveness to ERK signaling and marked developmental robustness in growing teleost skin.

## Introduction

Tissue morphogenesis requires coordination of cellular behaviors and integration of signaling. Multiple cellular processes, including cell proliferation and rearrangements, guide morphogenesis, but their relative contributions has been quantitatively dissected only in limited instances^1–4^. Moreover, the resilience of these morphogenetic processes to perturbations in cellular dynamics remains largely unexplored. This is particularly relevant in rapidly growing and elongating tissues during development, as morphogenetic processes would be tightly coordinated across tissues^5,6^. Here, we address the cellular processes and molecular mechanisms that govern the robust elongation of the zebrafish epidermis.

The epidermis tightly envelops and protects underlying structures. During development, the vertebrate embryo preferentially elongates along the anterior-posterior axis, with the epidermis consequently undergoing directional expansion to accommodate this rapid growth. The mammalian epidermis is specified as a single layered epithelium, derived from the ectoderm, which then transits through a bilayer phase and progresses to become a multilayered and eventually keratinized structure^7–9^. The bilayered epidermis comprises of two epithelial cell layers: a basal stem cell layer and outer periderm layer, connected by desmosomes. Studies of vertebrate epidermis have shown that temporal alteration of the cleavage plane in basal epithelial cells, increased proliferation of basal and suprabasal layers and subsequent delamination of cells from basal layer dictates stratification and expansion of basal epithelial layer^7,9–13^. In contrast, the cellular and molecular mechanisms regulating rapid expansion of periderm during developmental growth are less well-studied and understood owing to the temporary nature of this tissue as it is shed *in utero*. Periderm functions as a transient mechanical barrier and protects developing basal stem cell layer until the epidermis matures, preventing abnormal fusion between adjacent epithelial tissues^14,15^. Further, gene disruptions related to periderm function or formation result in birth defects marked by interepithelial fusion, known as peridermopathies, underscoring the crucial role of the periderm in embryonic development^8^.

Zebrafish embryos are transparent and develop externally, yielding an opportunity to visualize complex biological processes at high spatial and temporal resolution. In zebrafish, the periderm is specified on the first day of life and is derived from the enveloping layer (EVL)^16^. In contrast, the basal stem cell layer is specified subsequently from non-neuralized ectoderm^17,18^. The epidermis remains a bilayer until metamorphosis, i.e. for about 20 days post fertilization, at which point it initiates further stratification^18–20^. As the epidermis maintains a bilayer structure for an extended duration in zebrafish embryos^21^, they present an ideal model system to investigate the cellular behaviors and signaling dynamics involved in periderm growth.

Directional growth of epithelia could be attained by a combination of cell intercalations driving neighbor exchanges, oriented cell divisions, increases in cell size, cell shape changes/reorientation, and/or addition of new cells from underlying tissues^22^. Autoradiographic studies show presence of proliferative cells in both periderm and basal layer of mouse limb buds cultured *in vitro*, suggesting that periderm could grow independently of basal layer^23^. It remains uncertain which cellular behaviors drive the directional growth of periderm and whether cells are exchanged between the two layers during such rapid phases of expansion. Further as epithelial tissues can alter their growth in response to mechanical forces exerted by adjacent embryonic tissues^24–26^, it is unclear how periderm cells respond to the mechanical challenge of rapid axial elongation while being tethered to the basal layer. Additionally, how periderm adjusts its growth when this mechanical challenge ceases is yet to be determined.

The Mitogen-activated protein kinase (MAPK) family of proteins transduce extracellular signaling to control cellular responses. Extracellular signal-regulated kinase (ERK) signaling has been implicated in regulating cell division, migration, differentiation, and growth - cell behaviors that are central to morphogenesis of epithelia^27,28^. In cultured mammalian keratinocytes, the exit from basal stem cell compartment is characterized by increase in pulsatile ERK activity^29^. Further, spatial propagation of ERK activity coincides with increased epidermal proliferation within basal epidermal layer of adult mice^30^. Axial elongation provides a herculean challenge to the nascent periderm, and it is unclear how early epidermal structures like periderm adjudicate signaling to regulate their cellular behaviors which contribute towards rapid directional growth.

Tissue growth is determined by an increase in total cell mass - which can be accomplished by an increase in cell size and/or cell numbers^31^. Upon perturbations, some tissues have limited plasticity, such as pancreatic tissues, as their growth is constrained by the initial pool of progenitor cells. In contrast, tissues like the liver are not bound by cell numbers and can robustly achieve their target size through hypertrophy^32^. Although specific cell behaviors can drive periderm growth, it is unclear whether periderm maintains flexibility in employing these cell behaviors upon perturbations. The degree of resilience within the epidermal cells and the potential impact of disruptions on robustness of epidermal growth remains unclear.

Here, we established a set of tools to quantitatively describe *in toto* signaling dynamics and cellular behaviors during growth of the zebrafish periderm. Our findings highlight developmental changes in responsiveness of periderm cells to ERK signaling during animal growth and indicate an underlying robustness in epidermal cells in overcoming roadblocks to division.

## Results

### Periderm growth occurs by oriented cell divisions

Axial elongation in zebrafish embryos begins post gastrulation, peaking at 24-48 hours post fertilization (hpf) and continues albeit at slower rates as the larvae grows into adults. Post 24 hours post fertilization (hpf), this coincides with a rapid expansion of the embryonic periderm. As the embryo undergoes a disproportionate increase in length compared to its width, the periderm must similarly grow to accommodate this elongation. To understand the cellular mechanisms that drive rapid periderm expansion, we developed transgenic *occludinb-GFP* and *krt4:*H2A-mCherry zebrafish lines, enabling the labeling of cell membranes and nuclei in the periderm (Figure 1A-B, S1A). Employing an imaging setup that allows unobstructed posterior elongation, we conducted live imaging of periderm growth for 12 to 14 hours, starting at 28 to 30 hours post-fertilization (hpf) (Figure 1A-B). Computational segmentation of individual cells facilitated the quantification of various cell attributes, for example cell size and shape, and behaviors, including neighbor exchanges and cell divisions. Axis elongation driven by convergence extension, has been extensively studied in many systems, including embryonic ectoderm of *Drosophila melanogaster*^33^*, Xenopus laevis* gastrulation^34^, zebrafish^35^ and mouse tissues^36,37^. In these systems, the epithelium becomes narrower and longer over time by cell intercalation through directional exchange of neighbors. In contrast to these tissues, our observations of neighbor exchange events between periderm cells during the rapid expansion phase were minimal (Figure 1C), with only 0.8% of cells undergoing such exchanges (Figure S1B). Notably, we observed 27% of *Drosophila* cells undergoing neighbor exchange during germ band extension^38^, validating the efficiency of our computational pipeline to automatically detect neighbor exchanges (Figure S1B- C). The lack of neighbor exchanges in this tissue argues that neither convergent extension nor cell addition from the basal layer drives the rapid extension of the periderm.

**Figure 1:**
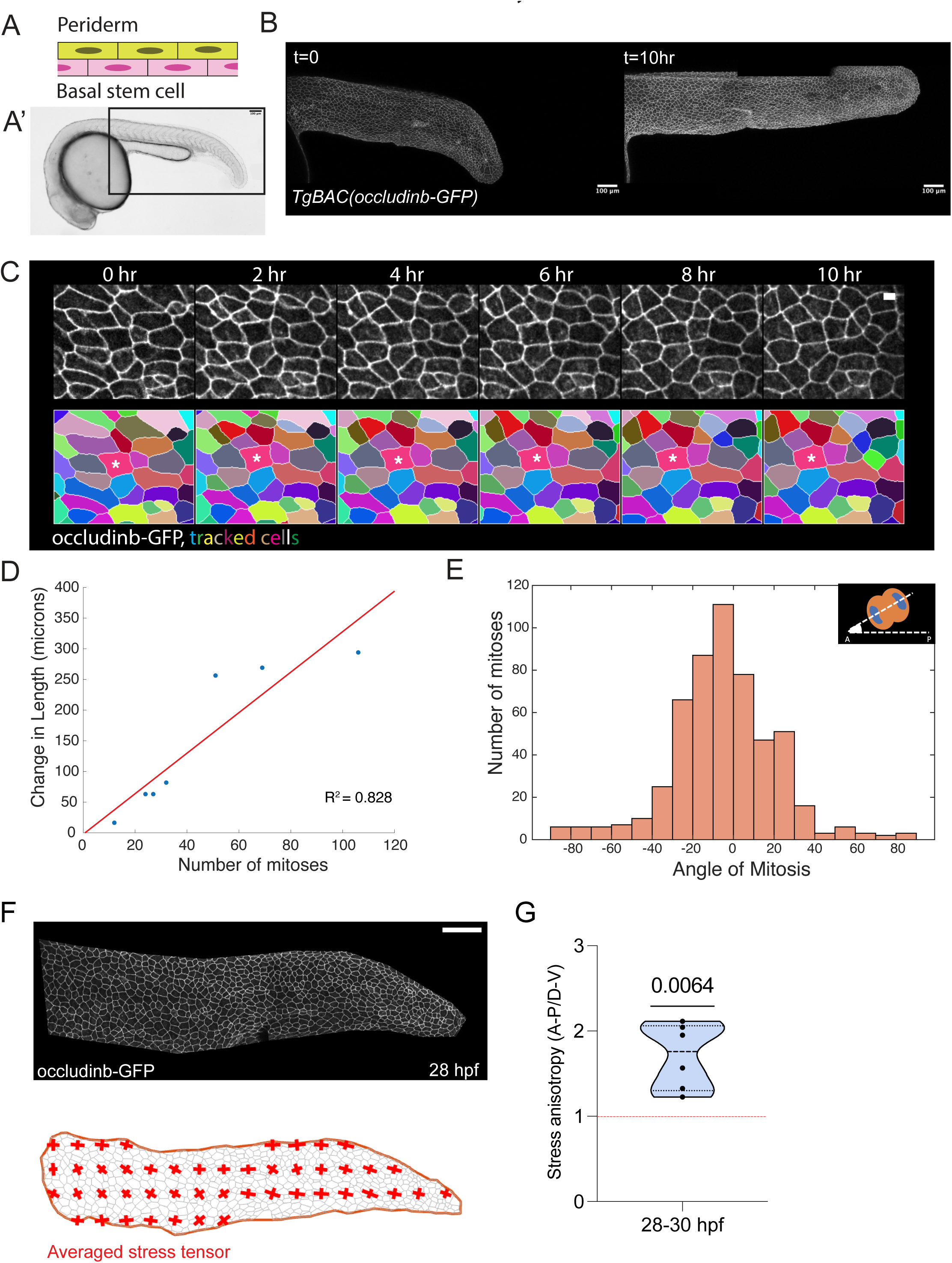
Rapid periderm expansion is facilitated by oriented cell divisions. Zebrafish embryonic epidermis comprises of the outer periderm and inner basal stem cell layer (A). We imaged the expansion of periderm covering the posterior tail bud from 28 hpf onwards (A’). Scale bar – 100 µm. (B) Stills from timelapse movies of transgenic embryos expressing *occludinb*-GFP at 0 and 10 hours post imaging (hpi), showing the expansion of the periderm during the imaging. Scale bar – 100 µm. (C) Higher magnification stills from timelapse of periderm and the corresponding cells tracked, show the cells maintain their neighbors during imaging. Scale bar – 10µm. (D) Quantification of the increase in length of the embryo and the number of mitoses observed during that time, shows a high correlation with R^2^= 0.828, suggesting that embryo elongation is facilitated by cell divisions. N= 7 embryos. (E) Quantification of the angle of cell divisions, which is defined as the angle between the mitotic spindle and the anterior-posterior axis of the embryo (inset). Histogram shows that cell divisions are aligned along the A-P axis of the embryo. 0-being along A-P axis. N = 7 embryos. (F) Image of periderm tissue expressing occludinb-GFP at 28 hpf and corresponding average stress tensor computed from the still image of the embryo, shows higher stress along A-P axis compared to D-V axis. Scale bar – 100µm. (G) Ratio of the anisotropy of stress along A-P vs D-V for multiple embryos between 28-30 hpf, shows that this is significantly higher than 1. One-sample t-test p = 0.0064, N= 6 embryos

Intriguingly, we observed several mitotic events, prompting an investigation into their contribution to periderm elongation. Our analysis revealed a proportional relationship between the change in periderm length and the number of observed mitoses (Figure 1D), indicating the substantial contribution of mitosis to periderm expansion. Furthermore, the orientation of these cell divisions predominantly aligned with the anterior-posterior axis of the embryo, the primary axis of periderm growth (Figure 1E), suggesting that oriented cell divisions facilitate the anisotropic elongation of the tissue. While cells exhibited a moderate increase in size, as evidenced by the rise in average cell area (Figure S1D), the average aspect ratio decreased over time (Figure S1E). These findings suggest that while cell size increase contributes modestly to periderm expansion, it remains a secondary factor for early periderm growth. Collectively, our results underscore the predominant role of oriented cell divisions in driving the rapid elongation of the periderm.

### Periderm tissue experiences anisotropic stress during rapid axial elongation

The periderm undergoes significant elongation along the A-P axis while retaining an almost constant width. We hypothesized that differential tension, potentially due to the uneven growth of the underlying embryonic structures, might influence the orientation of cell divisions. To assess stress within the tissue, we attempted to perform laser ablations. In this technique, a cell-cell junction is ablated (often using a UV-laser) and a recoil is observed that is proportional to the tension experienced by that junction. However, we found that, unlike in other tissues^39,40^, we could not observe a proper recoil (data not shown), which we attributed to the dissipative ability of the stratified epidermis. Therefore, we employed a Bayesian force inference method^41,42^ to estimate tension at cell-cell junctions. By analyzing static images of the tissue with cell membranes labeled using *occludinb-GFP* (Figure 1F), we inferred tensions along individual cell edges. These tensions were then averaged locally across several cells to infer a map of stress in the tissue. Our analysis revealed that the periderm tissue experiences higher stress along its anterior-posterior (A-P) axis, which corresponds to its major axis of growth (Figure 1F, S1F), than dorsal-ventral (D-V) axis. We then estimated the anisotropy in stress along these two axes, with the A-P axis experiencing approximately 1.7 times greater stress (Figure 1G). This observation of anisotropy aligns with the distribution of cell orientations, wherein cell long axes are preferentially aligned along the A-P axis of the embryo (Figure S1G). Notably, this alignment persists as the embryo continues to elongate beyond 48 hpf, albeit at a slower rate. These observations support the computational inference of stress distribution and are consistent with a model in which predominant stress along the A-P axis directs cell orientation and influences the axis of cell division.

### ERK activity is predictive of cell proliferation in the periderm

To investigate which molecular signals might control cell proliferation, we focused first on MAPK signaling, which plays a pivotal role in many contexts of tissue growth^27,28^. To determine the significance of MAPK signaling in periderm growth, we treated embryos with a MEK inhibitor (PD0325901) and evaluated cell cycling. To delineate cycling cells, we generated a transgenic line expressing Venus-hGeminin, a marker for cells in the S-phase^43^, directed by the *krt4* promoter. Our analysis revealed a 32% and 45% reduction in cycling following MEK inhibition, evident at 6- and 18-hours post-treatment respectively (Figure 2A-B, S2A-B). These findings implicate MAPK signaling as a regulator of peridermal cell proliferation.

**Figure 2:**
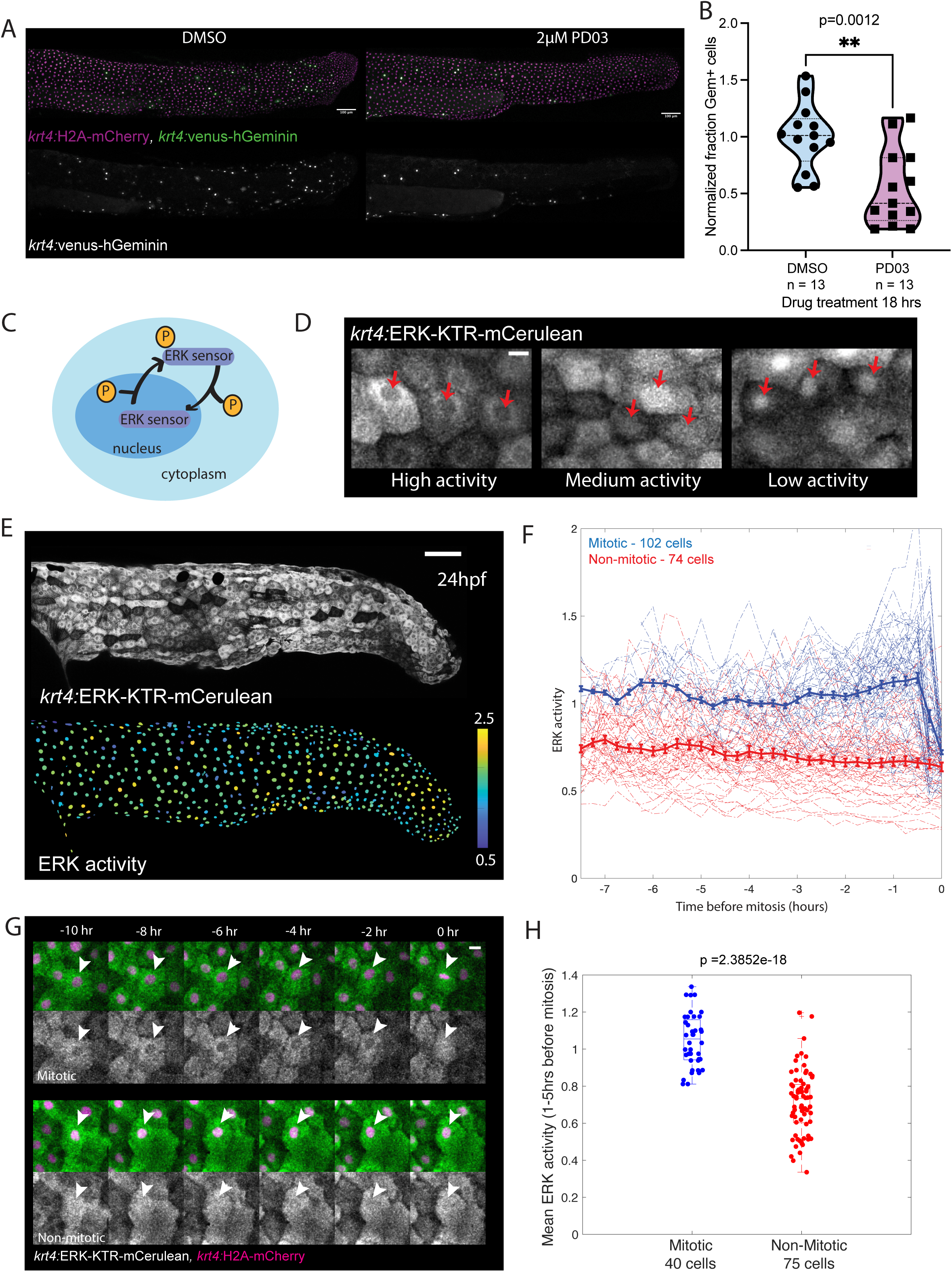
ERK activity is predictive of cell proliferation in the periderm. (A) Stills from transgenic embryos expressing *krt4:*H2A-mCherry *and krt4*:venus-hGeminin treated with DMSO or 2µM PD0325901 for 18hrs. Embryos treated with PD03 have significantly reduced proliferation seen by decrease in geminin positive cells. Scale bar – 100µm. (B) Quantification of fraction of geminin positive cells in embryos treated with DMSO or PD0325901, shows that PD03 treated embryos have significantly decreased proliferation as seen by reduction in normalized fraction of geminin positive cells. Average of DMSO = 1, n=13 embryos, Average PD03 = 0.5455, n=13 embryos. P=0.0012. This suggests that ERK activity is required for propagation through cell cycle. (C) Schematic of ERK translocation sensor. The sensor shuttles between nucleus and cytoplasm upon phosphorylation. The ratio of cytoplasmic to nuclear signal gives the inferred activity of ERK in single cells. (D) Examples of high, intermediate, and low ERK activity in periderm cells. Sensor shuttles into nucleus as ERK activity decreases. Scale bar – 10µm. (E) Snapshot of *krt4:*ERK-KTR-mCerulean embryo expressing the ERK sensor in the periderm and the corresponding heat map of ERK activity. ERK activity is high in most cells of the periderm without any specific spatial pattern at 28hpf. Scale bar – 100µm. (F) Quantification of ERK activity levels in single cells of the periderm. Cells were classified as mitotic (blue) if they divide during the imaging. Mitotic cells (blue) have higher ERK activity compared to non-mitotic cells (red), in the hours leading to division. T=0 is time of nuclear division. (G) Stills from timelapse of mitotic and non-mitotic cells showing higher ERK activity in mitotic cells. Arrow heads point to mitotic cell in upper panel and non-mitotic cell in lower panel. Scale bar – 10µm. (H) Average ERK activity 1-5 hours before mitosis is significantly higher than that compared with random 4 hours window for non-mitotic cells. Median of average ERK activity for mitotic cells – 1.054, n= 40 cells and non-mitotic cells – 0.734, n = 75 cells, t-test p = 2.3852e-18.

Live imaging of the embryonic skin has detected ERK activity in periderm^44^. Therefore, to define the dynamics of MAPK signaling in periderm, we generated transgenic fish expressing the ERK- KTR translocation sensor^45^ fluorescently tagged with mCerulean, under control of the *krt4* promoter. Using the ratio of nuclear to cytoplasmic signal, we can determine the signaling status of cells in the periderm (Figure 2C-D). During early zebrafish development i.e. <6 hpf, an ERK-KTR sensor was recently shown to be responsive to both ERK and Cdk activity^46^. Therefore, to establish the specificity of our sensor, we treated embryos with MEK inhibitor and Roscovitine, a Cdk inhibitor. We found that cells specifically reduced inferred ERK activity upon MEK inhibition when compared with Cdk inhibition (Figure S2C-D). This suggests that, in this tissue, ERK-KTR sensor responds mainly to MAPK and it is only affected to a limited extent if any by Cdk activity.

We observed no stereotyped spatial pattern of ERK activity at 28 hpf (Figure 2E). To determine how ERK activity is coupled with cell behaviors in the periderm, we conducted a longitudinal analysis, monitoring ERK activity at the single-cell level over time. As periderm cells mainly undergo proliferation, we measured the relationship of MAPK signaling to cell cycling. We monitored ERK activity dynamics in periderm cells and classified cells as mitotic and non-mitotic based on their ability to divide during 12 hours of the imaging experiment. We found that mitotic cells have higher average ERK signaling when compared with non-mitotic cells, especially in the 1 to 5 hours leading to up to division (Figure 2F-H). Further, using logistic regression, we find that average ERK activity in this 4-hour window could predict with 84% accuracy the ability of a cell to undergo mitosis (Figure S2E). This combined with our results on MAPK inhibition indicate that ERK activity is coupled to cell proliferation and is instructive for cell division.

### ERK is sufficient for S-phase entry of periderm cells during rapid axial elongation

Monitoring single-cell dynamics in MCF10A cells has shown that high ERK activity can be predictive of cell divisions^47^, however, this connection has not been demonstrated in an *in vivo* vertebrate setting. To test whether ERK activity is sufficient for proliferation when periderm undergoes rapid expansion, we established an optogenetic system to activate ERK in a spatial and temporally controlled manner. This system is based on the ability of the photo-convertible protein Dronpa to dimerize and sterically inhibit enzymes when inserted at both their N- and C-termini (Figure S3A). To control ERK activity in the periderm, we expressed under the direction of *krt4* promoter, a photo switchable phospho MEK E203K whose activity can be reversibly regulated by light, inactive at 405 nm and active at 488 nm (Figure S3A)^48,49^ .

Since ERK is active in many cells in the periderm, to rigorously test the ability of the optogenetic system to activate ERK, it is necessary to block endogenous ERK signaling. Therefore, we first examined candidate upstream ligand-receptor interactions that modulate ERK signaling at this stage. Treating larvae with EGFR inhibitor results in decreased pERK levels observed in the epidermis via western blot analysis^50^ and FGF has been implicated in axis elongation^51^. Using compounds that specifically block these receptors, we found that while FGFR inhibition with BGJ398 had moderate inhibitory effects on ERK signaling in periderm cells, EGFR inhibition with PD168393 had much stronger effects (Figure S3B-C). This suggests that EGF is the prominent ligand regulating ERK signaling in the tissue at this stage. To determine whether the optogenetic platform could stimulate ERK signaling in the periderm, we combined this line with our ERK-KTR sensor and treated the embryos from 18 to 24 hpf with EGFR inhibitor, PD168393 (Figure 3A). This led to a complete block in endogenous ERK activity as seen in Figure S3D-E, 3C. Upon Dronpa photoconversion to the dark state for 10 min, we observed both a drop in fluorescence (likely implying reduced Dronpa dimerization, see Figure 3B) and about a 70% increase in average ERK activity visualized by translocation of ERK-KTR sensor (Figure 3C-D). Conversely, in regions where Dronpa was not photoconverted, we observed 0.27% decrease, i.e. no significant change in ERK activity levels during the same timeframe (Figure 3C-D, S3D-E). These experiments validate the use of the optogenetic construct in periderm cells.

**Figure 3:**
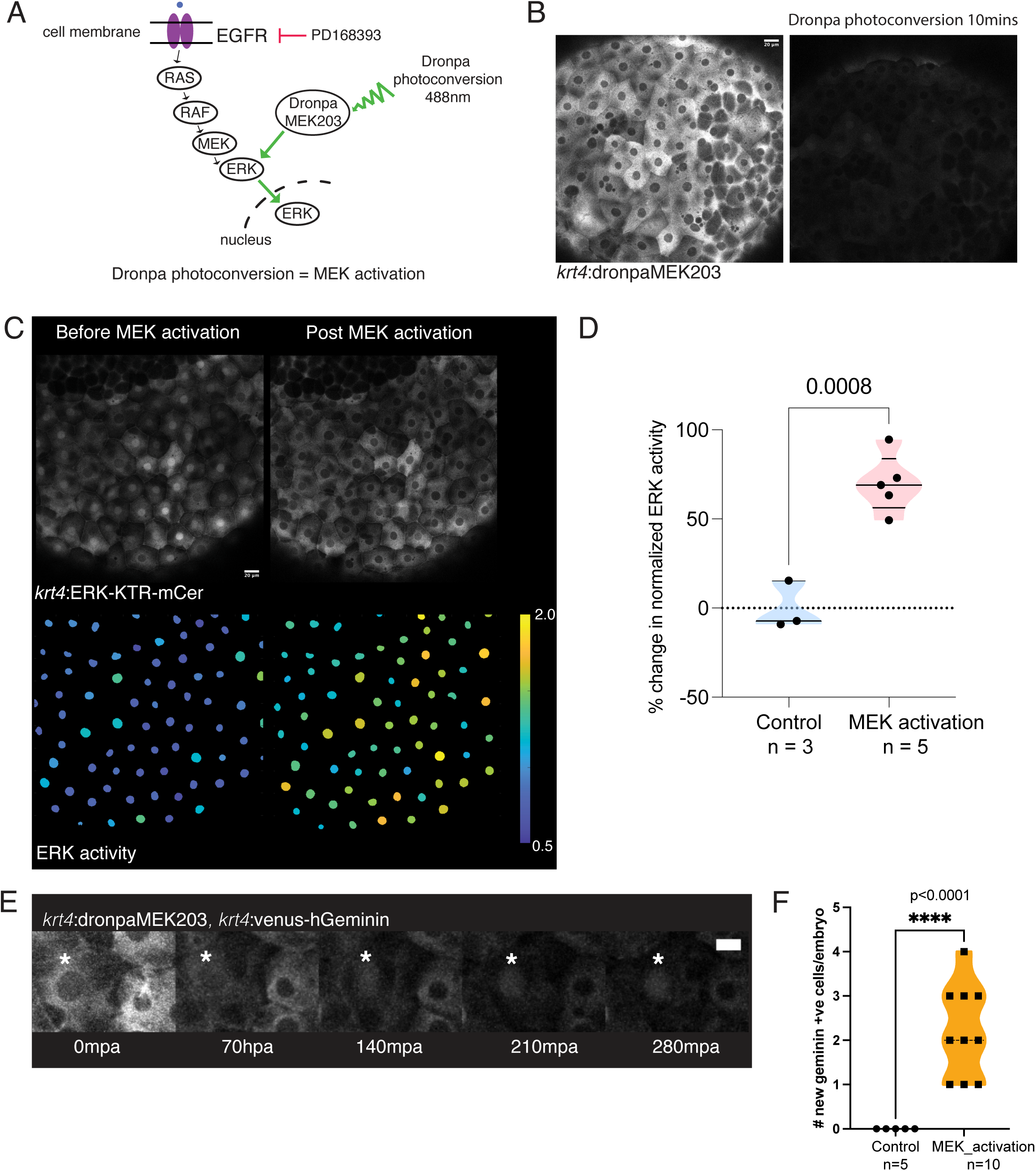
Optogenetic induction of ERK activity promotes S-phase entry in periderm cells during rapid expansion. (A) Scheme of experimental set up for turning off endogenous ERK signaling and ensuring that ERK mediated signaling is through Dronpa photoconversion. (B) Maximum z-projection of transgenic embryos expressing *krt4:*Dronpa-MEK203. Strong illumination with laser at 488nm for 10mins leads to photoconversion of Dronpa, leading to loss of its fluorescence. Scale bar – 20µm. (C) Stills from representative embryos expressing Dronpa-MEK203 and ERK-KTR-mCerulean, showing localization of ERK sensor and corresponding ERK activities. Before MEK activation, sensor shows nuclear localization, i.e. low ERK activity as embryos are in EGFR inhibitor. However, upon MEK activation (Dronpa photoconversion), sensor dramatically translocates to the cytoplasm indicating high ERK activity. Scale bar – 20µm. (D) Quantification of change in normalized ERK activity upon MEK activation or no activation, shows 69.88% increase in normalized ERK activity upon MEK activation. Upon no activation the change in normalized ERK activity was -0.27 %. Unpaired t-test, p = 0.0008 (E) Still from timelapse of representative embryonic periderm cells expressing *krt4:*Venus-hGeminin and *krt4:*DronpaMEK203. Geminin expressed in the nucleus becomes active as the cytoplasmic Dronpa fluorescence diminishes, i.e. MEK is activated using optogenetics. * indicates cell activating geminin expression. Scale bar – 10µm. (F) Quantification of number of cells entering S-phase as seen by turning on geminin expression. As the embryos are in EGFR inhibitor for 5-6 hours before the experiment, no new cells enter S-phase during the imaging. In contrast, with MEK activation, a small number of cells enters S-phase. Control average = 0, MEK activation average = 2.2 cells/embryo. Welch’s t test p<0.0001.

To determine whether this spatially controlled ERK activity can induce proliferative responses in the periderm, we combined our optogenetic lines with *krt4:*Venus-hGeminin as expression of geminin reflects cells entering S-phase. We photoconverted Dronpa and visualized cycling cells using the *krt4*-directed Venus-hGeminin reporter. As the embryos are in EGFR inhibitor, we found that no new cells enter S-phase during our imaging (Figure 3F). However, experimental activation of ERK led to a small number of cells entering S-phase within 3-4 hours after stimulation (Figure 3E-F). The moderate potency of this response could be ascribed to the necessity for prolonged EGFR-mediated signaling and/or to additional EGFR-dependent signals that are not activated by MEK. To corroborate this, our logistic regression yielded an AUC of 0.84 (Figure S2E), while MEK inhibition only blocked about half of the cells from dividing (Figure 2A-B, S2A-B). Collectively, these observations suggest that ERK activity is an efficient readout of EGFR signaling and a potent activator of the cell cycle, but that additional factors in the EGFR cascade contribute to mitosis. Consistent with this, we observed a notably greater suppression of proliferation with the EGFR inhibitor when compared with MEK inhibition (Figure S3F-G). Nevertheless, our results suggest that ERK activation is sufficient to increase S-phase entry in periderm cells, although additional signals from continued EGFR signaling are ostensibly required during rapid periderm expansion.

### Periderm proliferation decreases and ERK activity becomes pulsatile as elongation rate slows

The rate of embryo elongation slows after 48 hpf to approximately half the rate of expansion observed during the initial day (Figure S4A-B). To assess the cell cycle status in greater detail and simultaneously examine ERK signaling, we developed a transgenic line expressing a cell cycle translocation sensor Cdk2 KTR^52^ tagged with mCherry, under the *krt4* promoter. By evaluating the ratio of cytoplasmic to nuclear signal, we were able to quantify Cdk activity (and thus the cell cycle status) in individual cells within the periderm tissue (Figure 4A). Additionally, we generated another transgenic line expressing H2A-iRFP (far red) for nuclear labeling under the *krt4* promoter. To test the responsiveness of our sensor to Cdk activity, we treated embryos with the Cdk inhibitor Roscovitine and quantified the number of mitotic cells. Following a 4-hour treatment, our observations revealed a reduced proportion of cells exhibiting high sensor activity, with 32.6% in the DMSO-treated group vs only 21.9% in the Roscovitine-treated group, i.e., 33% reduction, indicating the sensitivity of the sensor to Cdk activity (Figure S5A-B). The moderate response could be attributed to lower drug concentration used in concert with shorter incubation times. Leveraging these lines, we tracked Cdk activity in both mitotic and non-mitotic cells, revealing expectedly high Cdk activity in mitotic cells and markedly lower activity in non-mitotic G1 cells (Figure 4B-C). Employing these transgenic lines, we detected a decline in the number of mitotic cells from 44% at 28-30 hpf compared to 5% at 52-54hpf, (Figure 4D-E). Consistent findings were obtained through analysis of the geminin reporter line, with 36% at 28 hpf and 6% at 52 hpf (Figure S5C-D). These observations strongly indicate a reduction in cellular proliferation concurrent with the slowing pace of embryo growth.

**Figure 4:**
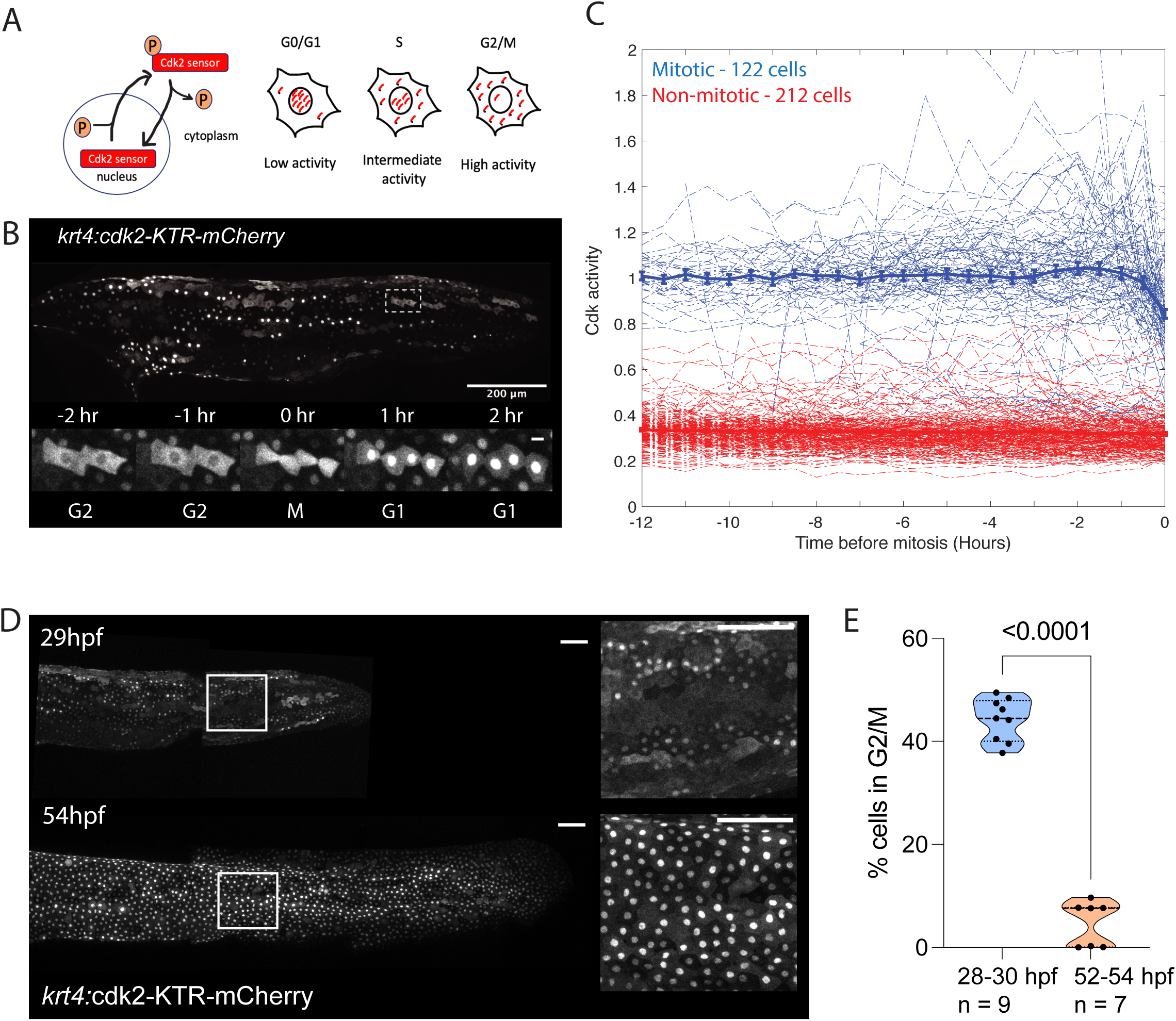
Periderm proliferation decreases as axial elongation rate slows. (A) Schematic of working of Cdk2 sensor. The fluorescently tagged protein shuttles in-and-out of the nucleus. Based on the cytoplasm to nuclei signal ratio, we determine the cell cycle status. (B) Maximum projection view of representative embryo expressing *krt4:*Cdk2-KTR-mCherry, scale bar – 200µm. Higher magnification view of cells in boxed region of the embryo tracked through time, shows translocation of sensor from cytoplasm to nucleus as cells transition from G2/M to G1 phase post-mitosis. Scale bar – 10 µm. (C) Quantification of Cdk activity in periderm cells classified as mitotic or non-mitotic based on their ability to divide during the imaging. Expectedly mitotic cells have much higher Cdk activity compared to non-mitotic cells. (D) Maximum projection view of posterior region of embryos expressing *krt4:*Cdk2-KTR-mCherry at 29 hpf and 54 hpf. The localization of the tagged sensor shifts predominantly towards the nucleus in the later stages, signifying the entry of cells into the G1 phase. Higher magnification view of boxed regions showing the sensor translocation. Scale bar – 100 µm (E) Quantification of percentage of mitotic cells at 28-30hpf and 52-54 hpf, shows significant reduction in the number of cells in G2/M at the latter stage. Average percentage of mitotic cells at 28-30 hpf – 44.23%, n = 9 embryos; at 52-54 hpf – 4.72%, n = 7 embryos. Welch’s t test p <0.0001

We continued to monitor ERK activity dynamics during later phases of reduced growth rates, finding a more pulsatile nature of ERK signaling during 54-74 hpf as compared to 30-50 hpf (Figure 5A). To quantify and re-assess this observation, we used the continuous wavelet transform (CWT)^53^ to estimate the presence of pulses of given duration at specific times. Our findings indicate 34% increase in pulses of ERK activity lasting between one and three hours at 54-74 hpf compared to 30-50 hpf, as evidenced by the magnitude of the ERK CWT values during 54-74 hpf (Figure 5B-C). Moreover, these pulses were asynchronous, without a defined temporal pattern, as indicated by the dispersed distribution of high activity across the plot (Figure 5B). Thus, in contrast to the period of rapid embryo expansion at 30-50 hpf and relatively stable ERK activity, ERK signaling dynamics at 54-74 hpf demonstrated increased variability and pulsatility concomitant with reduced cell division and axial growth.

**Figure 5:**
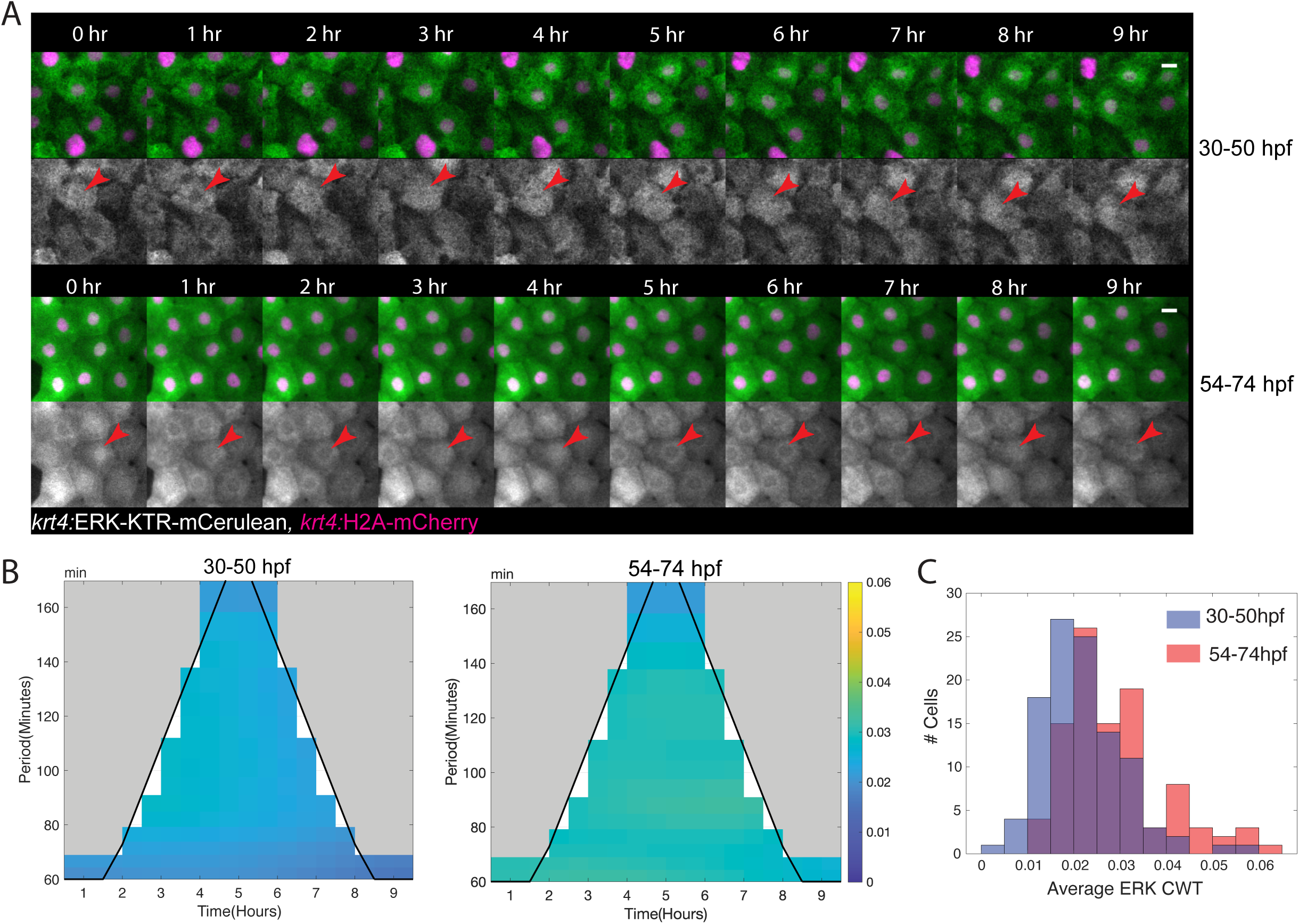
ERK activity becomes more pulsatile as periderm expansion slows. (A) Stills from timelapse imaging of representative periderm cells expressing *krt4:*ERK-KTR-mCerulean and *krt4:*H2A-mCherry. While some pulses of ERK activity are seen at 30-50 hpf (top panel), ERK activity becomes increasingly pulsatile from 54-74 hpf (bottom panel). Scale bar – 10µm. (B) Mean of continuous wavelet transform of ERK activity signal for 30-50 hpf embryos during randomly chosen 5-hour time window of non-mitotic cells, averaged across the individual CWT plots for individual cell ERK traces. Traces derived from 5 separate embryos, with total 107 cells (traces). For 54-74 hpf embryos, cells from 4 separate embryos used with total 99 cells (traces). The plots were modified to exclude CWT results along low-confidence regions of the trace which are confounded by edge-effects. There is a statistically significant difference in the mean of the CWT plots between 30-50 hpf and 54-74 hpf (C) based on the one-sided KS test, with a p-value of 4e-5, suggesting higher average CWT value at 54-74 hpf implying more pulsatile ERK activity.

### Shift in peridermal responses to ERK activity

The observed changes in ERK signaling dynamics coupled with the concurrent decrease in proliferation indicators prompted us to investigate whether cellular responses to ERK signaling are altered as embryo growth decelerates. To achieve this, we used a multiplex approach to image both Cdk and ERK activities via a combination of Cdk2-KTR-mCherry and H2A-iRFP to track cell cycle status, along with ERK-KTR-mCerulean to monitor ERK activity, in large numbers of individual periderm cells. We then used a threshold of ERK activity to study the cell cycle status of cells experiencing high ERK activity. Intriguingly, we discovered the existence of two distinct cell populations: a population with high ERK and high Cdk activities and a distinct population of cells characterized by high ERK but low Cdk activity. The former population is indicative of cells with elevated ERK signaling undergoing mitosis while the latter comprises cells in G1 phase that are responding to ERK signaling. On tracking cells from 30-50 hpf we found that approximately 21% of total cells belong to the former population, while 14% of total cells belonged to the latter population (Figure S6A).

To track the dynamics of these cell populations over time, we analyzed cell behaviors in 5-hour windows and defined the prevalence of these populations across different time points (Figure 6A). Our analysis revealed a decline in the percentage of high ERK cells exhibiting high Cdk activity from 71% to 33% over time, indicating a reduction in proliferative responsiveness to ERK signaling (Figure 6A). Conversely, the population of cells with high ERK but low Cdk activity significantly increased over time, rising from 28% at 40-45 hpf to approximately 66% at 55-60 hpf (Figure 6A). Further examination revealed that, as expected, post-division Cdk activity decreased as cells entered the G1 phase (Figure 6B, D). In contrast, ERK activity exhibited sporadic patterns of activation and deactivation, contributing to the heterogeneity in cell responses to ERK signaling (Figure 6C-D). There is a possibility that cells exhibiting high ERK activity may eventually activate Cdk activity. However, since we were unable to track cells beyond 20 hours, we cannot definitively establish this. Collectively, our data suggest a shift in cellular responsiveness to ERK signaling following mitosis.

**Figure 6:**
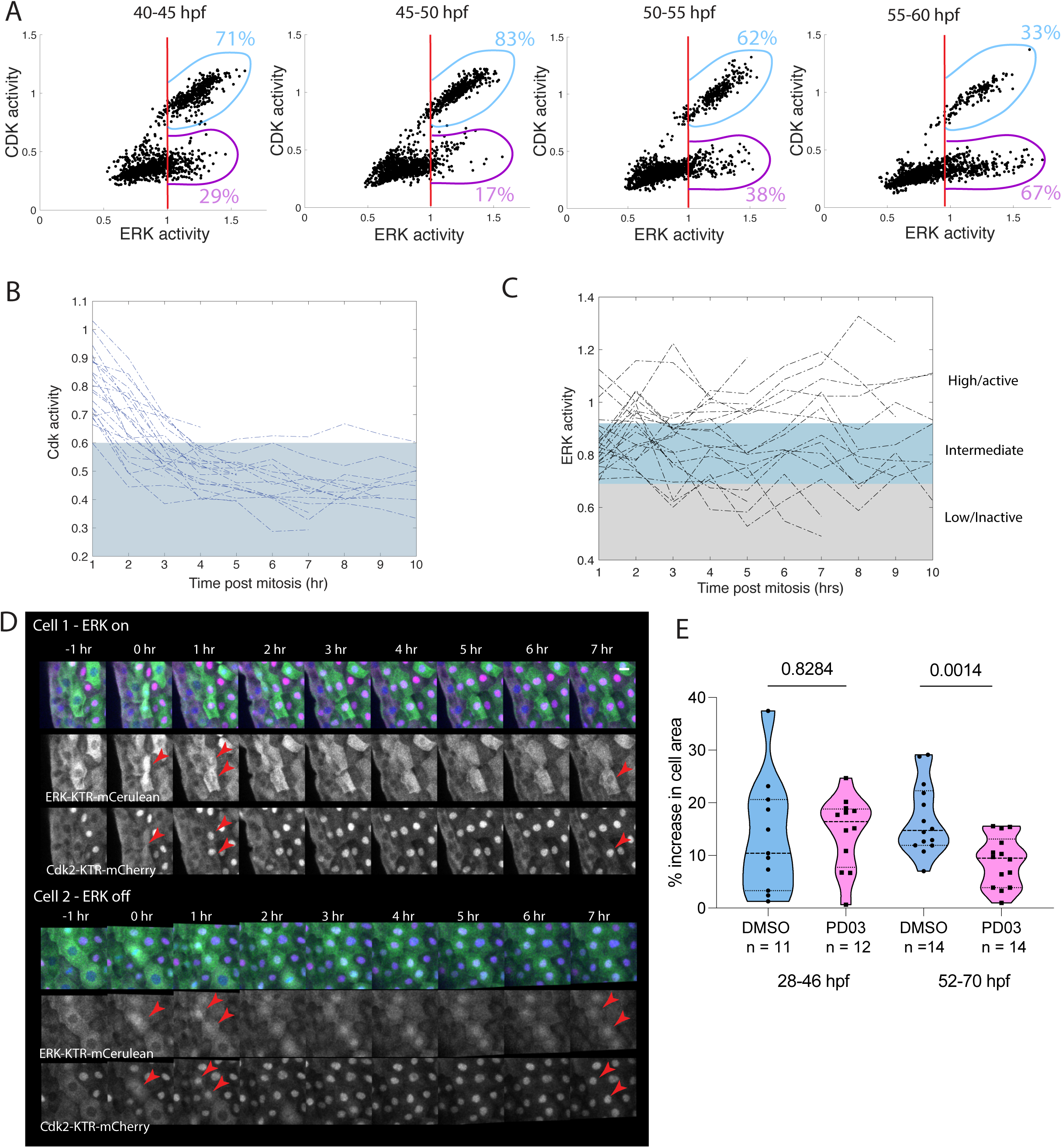
Cells switch from proliferative to non-proliferative responses to ERK signaling as periderm expansion slows. (A) Scatter plot of ERK and Cdk activities in single cells of the periderm at respective time windows. The percentage of cells with high ERK activity and high Cdk activity (blue population), decreases over time, while that of high ERK activity and low Cdk activity (pink population) increases over time. There is a significant increase in the probability of being in the high ERK low Cdk group at later time points, computed using binomial distribution. p = 1.2817e-53. High ERK is denoted by cells with ERK activity >=1. N = 3 embryos, 210 cells. (B) Quantification and tracking of Cdk activity post cell division, shows that cells enter G1 following mitosis as seen by the drop in Cdk activity post mitosis. Grey shaded region represents regime of Cdk activity where cells are in G1. Time t = 0hr is nuclear division, data is plotted from 1 hr post-cell division. N = 4 embryos. (C) Quantification and tracking of ERK activity show that following cell division, cells sporadically inactivate ERK signaling as seen by the spread of ERK activity from 1 – 10 hr post mitosis. This leads to a more heterogenous population. Grey and blue shaded regions represent regime of inactive and intermediate ERK signaling based on PD0325901 drug treatment experiments. N = 4 embryos. (D) Maximum projection of representative periderm cells from embryos expressing *krt4:*ERK-KTR-mCerulean, *krt4:*Cdk2-KTR-mCherry and *krt4:*H2A-iRFP. Top panel follows a cell where ERK activity is on following mitosis, while bottom panel follows a cell with ERK activity turned off following mitosis. In both cases, Cdk activity drops dramatically following division, indicating cells in G1 phase. Scale bar – 10 µm. (E) Quantification of increase in average cell area upon DMSO or 2µM PD0325901 treatment for 18 hours with treatment beginning at 28 hpf or 52 hpf. While there is no significant change in the increase in cell area between control DMSO and PD03 treated embryos at 28 hpf, there is significantly less increase in area upon PD03 treatment at 52 hpf. Percentage increase in cell area at 28 hpf - DMSO = 13.56%, n =11 and PD03 = 14.41%, n = 12, p = 0.8284 Welch’s t test. Percentage increase in cell area at 52 hpf - DMSO = 16.86%, n = 14 and PD03 = 8.823%, n = 14, p = 0.0014 Welch’s t test.

The increase in pulsatile nature of ERK activity and non-proliferative behavior of cells led us to hypothesize that cells may exhibit modified responses to ERK inhibition during this stage. To investigate this, we administered a MEK inhibitor to embryos for 18 hours at either 28-30 hpf or 52-54 hpf, measuring the average change in cell size following the corresponding treatments. Periderm cells increase their area by 16.8% on average from 52 to 70 hpf (Figure 6E). Upon MEK inhibition, there was a ∼two-fold reduction in this increase, i.e. cells increased their area by only 8.8%. This suggests that ERK activity promotes cell hypertrophy at this later stage of periderm expansion. MEK inhibition did not affect cell area changes during the earlier stage of periderm elongation (Figure 6E). A possible interpretation of these findings is that ERK signaling is crucial for promoting both cell size increase and cell division from 28 to 46 hpf. Here, the cell hypertrophy would not be detected as cells with high ERK signaling would eventually divide to yield daughter cells with smaller areas. Subsequently from 52 hpf onwards, the primary role of ERK is to drive an increase in cell size when proliferation decreases, and the elongation rate slows down. This suggests that as elongation phase slows down, ERK activity promotes hypertrophic growth of cells, which contributes to tissue elongation.

### Periderm growth exhibits adaptive robustness

We have documented that the periderm mainly employs cell divisions to promote its rapid growth during axial expansion. To test the extent to which peridermal proliferation is required for axial elongation, we generated transgenic zebrafish with the *krt4* promoter directing expression of the cell cycle inhibitor p21^54^ with a downstream cleavable GFP tag. To examine periderm cells with p21 expression, we generated a transgenic line expressing Lifeact-Ruby under *krt4* promoter. Using the p21 line, we could experimentally reduce proliferation specifically in the periderm. Surprisingly, we observed that axial elongation occurred normally in these animals, as measured by rate of growth of the embryonic posterior between 28-46 hpf and 52-70 hpf (Figure S7A-B). Further, when periderm tissue was examined closely, we found that upon p21 expression, periderm cells had massively increased cell size, on average to about 4x time that of controls (Figure 7A-C) and at times reaching 8-10x size of control cells (Figure S7C). In addition to being bigger, periderm cell size showed greater variability as seen by the coefficient of variation in cell size upon p21 expression (Figure S7C). These findings indicate that periderm tissue employs hypertrophic cell growth to compensate for lack of proliferation in the tissue. Additionally, while some embryos were able to temper expression of the p21 transgene as assessed by GFP expression, presumably by epigenetic silencing, most maintained expression in majority of periderm cells and grew to viable adults. These p21-expressing adults had moderately increased cell size ∼1.3x times those of control animals (Figure 7D-F, S7D). The prolonged survival observed in these fish expressing p21 and the reduction in cell size and its variability seen in adults suggests potential compensatory mechanisms at work during development, such as an accelerated shift to basally derived periderm cells, that might resolve massive shifts toward hypertrophy as observed at earlier larval stages. Thus, periderm tissue can adapt to hypertrophy when proliferation is experimentally inhibited, revealing mechanisms of robustness in coordinating periderm and organismal growth over sustained periods.

**Figure 7:**
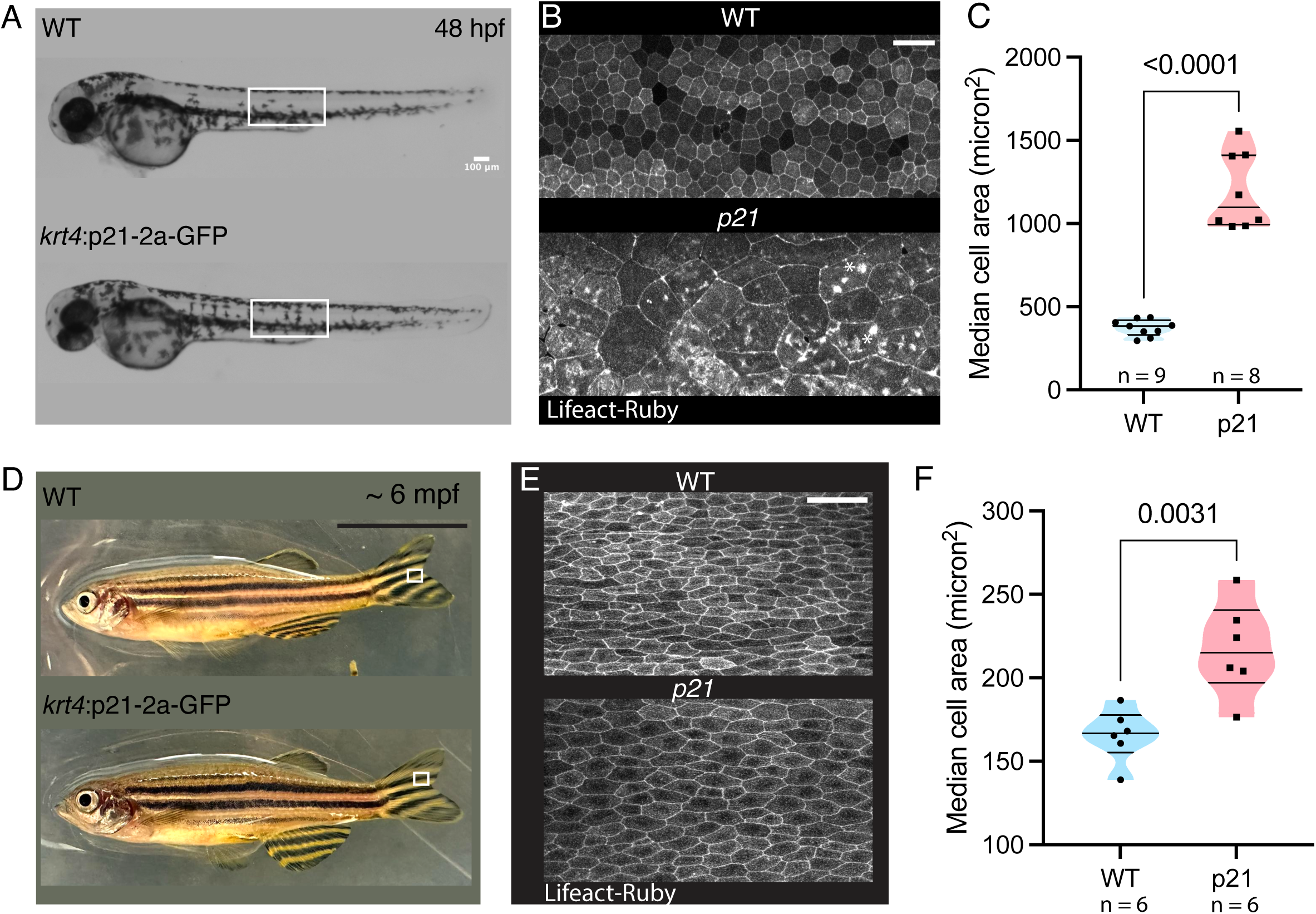
Periderm growth exhibits adaptive robustness. (A) Brightfield view of WT control and embryos expressing p21-2a-GFP under *krt4* promoter at 48 hpf, showing normal axial elongation. Scale bar – 100 micron. (B) Maximum projection view of periderm cells in the posterior trunk region of representative larvae boxed in A expressing Lifeact-Ruby under *krt4* promoter to outline cell boundaries. Periderm cell size is massively increased in embryos expressing p21. * shows abnormal accumulation of actin in periderm cells. (C) Median cell size quantified for control and p21 expressing embryos at 48 hpf. Average cell size in control embryos at 48 hpf, 373.1 micron^2^, n = 9; p21-expressing embryos, 1195 micron^2^, n = 8. P < 0.0001, unpaired t-test. (D) Brightfield view of WT and p21-expressing adult fish at 6 months, showing normal morphology. Scale bar – 1cm. (E) Maximum projection view of periderm cells in the fin region of representative adult fish boxed in D expressing Lifeact-Ruby under *krt4* promoter to outline cell boundaries. Periderm cell size is slightly increased in p21 expressing adults. (F) Median cell size of periderm cells covering fin region quantified in control and p21 expressing adults. Average cell size in control fish, 165.7 micron^2^, n = 6; p21-expressing fish, 217.2 micron^2^, n = 6. P = 0.0031, unpaired t-test.

## Discussion

### Oriented cell divisions in periderm growth

The periderm undergoes significant morphological changes in concert with embryo elongation. Here, we have dissected how tissue elongation is driven by underlying cellular behaviors and how these behaviors are linked to the growth factor signaling. Our data define a role for ERK signaling in regulating cell divisions and growth of periderm tissue during rapid axial elongation. We discover that periderm growth is primarily fueled by cell divisions aligned along the A-P axis, and notably, there is no observed cell intercalation or cellular exchange between the basal and periderm layers. As the rate of embryo elongation slows down, the burst in proliferation transits to hypertrophic cell growth of periderm cells (Figure 4D-E, 6E). Hence, we propose that during rapid expansion, when periderm experiences increased tension, signaling pathways upstream of ERK may trigger tissue proliferation, while stress anisotropy orients these divisions. Comparable to the spreading of zebrafish EVL^55,56^, these oriented cell divisions might serve to alleviate anisotropic tissue tension, potentially leading to subsequent decrease in proliferation in the periderm. Similar observations of bursts in proliferation followed by a decline, have been made in the periderm covering mouse upper limb buds in culture^23^, suggesting that this could be a conserved phenomenon.

### ERK activity pattern and cell cycle

By combining two transgenic KTR reporters, we determined the cell cycle status and ERK signaling in single cells of the periderm tissue covering the posterior region of zebrafish embryo. Using these new transgenic reporters, we identified a new population of cells with high ERK activity and low Cdk activity. Upon tracking these cells, we find that as cells undergo mitosis, they sporadically inactivate ERK, leading to a heterogenous population of post-mitotic cells. This corresponds with our observation of the increase in pulsatile nature of ERK activity in 54-74 hpf periderm cells (Figure 5). Pulsatile ERK activity, likely due to the transient activation of inhibitory pathways, has been noted across various tissue types during both proliferation and differentiation^29,57–59^. As the pulses observed in the periderm do not have a specific pattern, they could be a result of paracrine signaling or an influence of multiple ligands in the tissue. Cell cycle dependent ERK signaling dynamics have been implicated in fate specification and tissue differentiation in mammalian blastocyst ^60,61^. Hence, we hypothesize that cells that have exited mitosis and still exhibit elevated

ERK activity may represent cells in the initial stages of the differentiation, leading to the maturation of microridges and their interaction with the overlying glycans^62,63^. Further monitoring with markers for differentiation of the periderm and microridge maturation could help delineate these differentiating cells.

Intriguingly, we also observed that periderm cells respond differently towards inhibition of ERK activity at 52-70 hpf (Figure 6E). Notably, the inhibition of ERK activity at 52 hpf influences the augmentation in cell size. This observation aligns with our discovery that ERK activity and cell cycle status undergo alterations, indicating that differentiating cells may exhibit diverse responses to ERK signaling, likely involving hypertrophic growth^29,64^.

### Plasticity in the system

Our results suggest that periderm cells can switch from utilizing ERK signaling for proliferation and growth, toward directing hypertrophic growth at later stages. The transition from a proliferative to a non-proliferative state in cells as the growth rate diminishes has been previously documented in plants, fish scales, and mammalian cardiomyocytes^64–66^. We find that this transition appears to be adaptive; when proliferation is restricted, cells predominantly increase in size, occasionally growing to ten times or more their original size. This suggests that periderm cells are remarkably plastic in their ability to grow, ensuring that they maintain a protective barrier. Similar plasticity in the ability of cells to maintain an organ shape has been observed in *Drosophila* wing disc, a highly proliferative tissue. Upon inhibition of cell division specifically in the anterior region of the disc or within mutant clones, cells grew bigger, but this did not affect the growth or shape of the wing disc^67–69^. Remarkably, even the patterning of the wing disc was comparable to wild types. This suggests that tissue size and patterning are governed by total cell mass –a combination of cell size and cell number. Tissues like periderm and *Drosophila* wing disc retain the ability to flexibly increase total cell mass.

The ability of periderm cells to increase massively in size, could explain the rapid re-epithelialized of wounds in zebrafish larvae and adult epidermis^70^. Despite the normal axial elongation of the embryo, the p21 expressing larvae have abnormal actin accumulation in microridges (Figure 7B) and could therefore have altered skin properties, which might make them more susceptible to injuries. Further examination of stratification and skin properties could provide insights into potential alterations in these fish.

## MATERIALS AND METHODS

### Zebrafish

Wild-type and transgenic zebrafish of the Ekkwill (EK) strain aged between 3 to 15 months were selectively bred through timed mating for our experiments. Embryos were harvested in egg water and incubated at 28°C. Prior to experimentation, embryos were screened for the desired transgene expression. For imaging experiments, transgenic animals utilized were hemizygous, with clutch mates serving as internal controls. All procedures involving zebrafish were conducted in accordance with the ethical standards approved by the Institutional Animal Care and Use Committee at Duke University.

### Live imaging

Zebrafish embryos were anesthetized in 0.5X tricane. Alpha-Bungarotoxin (Invitrogen B1601) was injected into the embryonic heart of 24-26 hpf embryos, following which they were allowed to recuperate for an hour. After that embryos were screened for viability and immobilization by poking with hair loop and moved to imaging chamber with egg water. The imaging chamber was prepared using published mold^71^ with 1.5% agarose in 35mm glass bottom dish. Embryos were mounted laterally and once positioned were fixed in place with glass coverslip. Embryos were imaged in egg water on Leica SP8 and LAS X 2.01.14392 software with a HC FLUOTAR L 25x/0.95 W VISIR water immersion lens (Leica 15506374) at 0.75x zoom. As the posterior region is larger than the field of view of the microscope, multiple overlapping positions were acquired and stitched post-acquisition. Images were acquired at 1024X1024 resolution (0.606µm) and z- resolution of 1 micron. Fluorescent proteins were imaged using the following lasers – ERK-KTR-mCerulean – 458nm, occludinb-GFP- 488nm, venus-hGeminin – 488nm, DronpaMEK203- 488nm, Cdk-KTR-mCherry – 561nm, H2A-mCherry – 561nm, H2A-iRFP – 630nm, Lifeact-Ruby – 568nm.

### Pharmacological treatments

Embryos were screened at 24-26 hpf for appropriate transgene expression and equal numbers were placed in 40mm plates with agarose bed and treated with DMSO or drugs of appropriate concentrations. 4-6 hours or 18 hours post treatment, embryos were mounted laterally in the imaging chamber mold, immobilized with 1X tricane and imaged using laser scanning confocal. Following drugs were used – 2µM PD0325901 (MEK inhibitor), 2µM BGJ398 (pan-FGFR inhibitor), 50µM Roscovitine (Cdk inhibitor) and 10µM PD168393 (EGFR inhibitor).

### Optogenetic activation of ERK

Embryos were screened at 18-20 hpf for Dronpa expression and transferred to egg water containing 10µM EGFR inhibitor. Following 5-6 hours of inhibition, embryos were moved to imaging chamber and immobilized with 0.5X tricaine. Once positioned, they were fixed in place with glass coverslip and imaged in egg water containing 10µM EGFR inhibitor. For Dronpa photoconversion, embryos were exposed to 488nm laser by acquiring a z-stack continuously for 10 mins. To determine the efficacy of DronpaMEK203 to activate ERK signaling, following Dronpa photoconversion, image was acquired at 458nm for ERK sensor. For quantifying cell proliferation, embryos expressing venus-hGeminin and Dronpa were screened. While Dronpa was photoconverted using 488nm laser, Geminin image was acquired simultaneously using the same laser.

### Generation of transgenic fish lines

Following transgenic lines were used in this study.

|  |  |
| --- | --- |
| *Tg(occludinb-BAC-GFP)*<sup>pd1126</sup> | This paper |
| *Tg(krt4:H2A-mCherry)*<sup>pd400</sup> | This paper |
| *Tg(krt4:ERK-KTR-mCerulean)*<sup>pd401</sup> | This paper |
| *Tg(krt4:cdk2-KTR-mCherry)*<sup>pd402</sup> | This paper |
| *Tg(krt4:H2A-iRFP)*<sup>pd403</sup> | This paper |
| *Tg(krt4:Dronpa-MEK203)*<sup>pd404</sup> | This paper |
| *Tg(krt4:venus-hGeminin)*<sup>pd405</sup> | This paper |
| *Tg(krt4:Lifeact-Ruby)*<sup>pd406</sup> | This paper |
| *Tg(krt4:p21-2a-GFP)*<sup>pd407</sup> | This paper |

To generate the transgenic lines, we obtained pSKS plasmid containing *krt4* promoter sequence from Sagasti lab^72^. The digested plasmid was used as backbone for cloning the other components. The following primers were used –

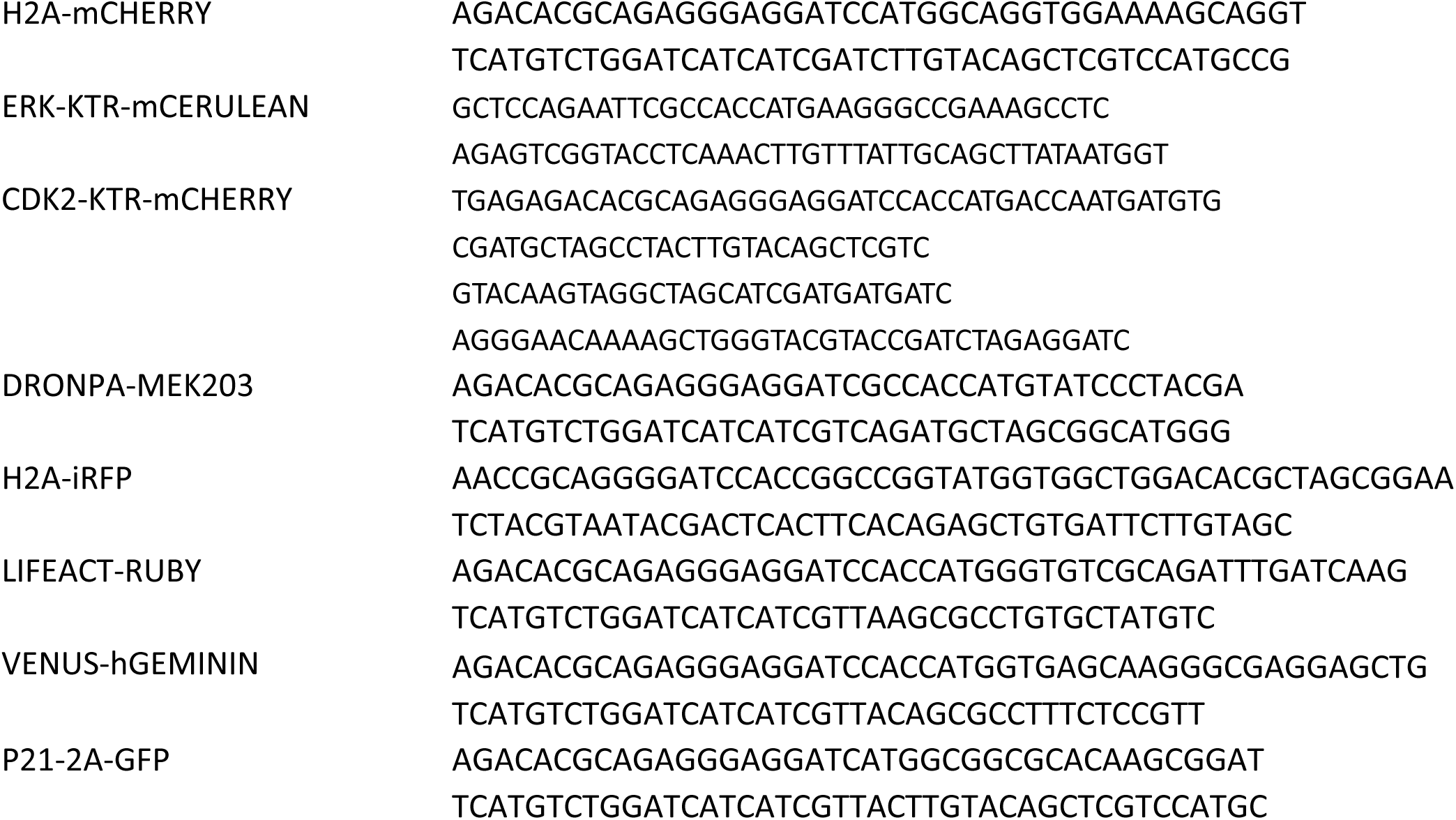

With the primers, the corresponding fragments were amplified from previously generated plasmids - H2A-mCherry^65^, ERK-KTR-mCerulean^65^, Cdk2-KTR-mCherry^52^, Dronpa-MEK-203^48^, Venus-hGeminin^65^ and Lifeact-Ruby. pTol2-UAS-zmiRFP was a gift from Edward Boyden. The corresponding pieces were cloned with PCR and inserted to make final construct using Gibson. The construct was co-injected with I-SceI into one-cell-stage EK embryos at a concentration of 100 ng/ml. Multiple founder lines were screened for each transgene and propagated.

### Generation of occludinB-GFP line

A bacterial artificial chromosome (BAC) containing *oclnb* CH211-214J8 (https://zfin.org/ZDB-BAC-050218-649) was modified as previously described^73^. For the C-terminal GFP fusion, targeting cassette was amplified from pBluescript-spGFP-FKF plasmid using following primers containing 50 bp *oclnb* homology sequences:

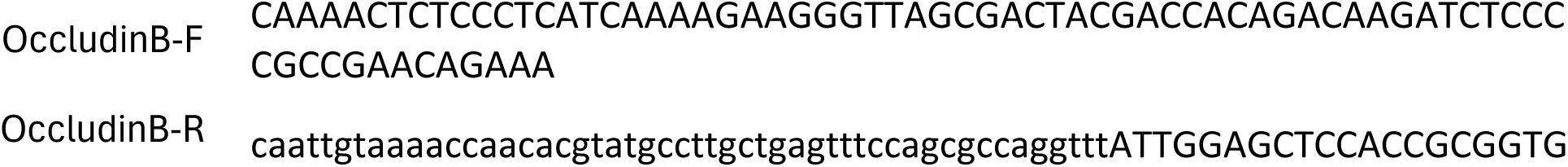

Homology arms were designed right before and after *oclnb* stop codon to replace the stop codon with the targeting cassette. FRT-KanR-FRT sequences were removed by flippase afterwards. For injection, *oclnb*-GFP BAC was linearized with PvuI enzyme (NEB), which cuts 110 kb upstream of occludin b gene coding region, and injected into one-cell stage embryos to generate *TgBAC(oclnb- GFP)pd1126*.

### Data analysis

Image processing was performed using Fiji, Cellpose, iLastik, Trackmate and custom written Matlab (MathWorks) R2022b software codes.

#### Cell tracking

Occludinb and H2A images for multiple positions were flattened and stitched using Fiji. Nuclei segmentation was carried out using iLastik. Annotated mitotic cells were manually checked and using custom matlab script we determined the angle of divisions. Occludinb-GFP images were segmented using Cellpose or EPySeg and the output was read by custom matlab script to determine cell properties like aspect ratio, orientation, area, etc. Using the segmentation data, cells were tracked with our custom matlab script. For neighbor exchange detection – cell outlines were dilated, and cell boundary subtracted to give a shell outline of cells. The neighbors with cell area in the shell were identified in subsequent frames. A neighbor change was marked if the neighbor was added or removed consistently for 4 consecutive frames proceeding the initial detection.

#### Bayesian force inference

Occludinb images acquired from multiple positions along the embryo were flattened and stitched in Fiji. Images were manually cleaned to remove underlying epidermis or other autofluorescence. The segmented outline from cellpose was fed into tissue analyzer. Following correction of tracks and segmentation, cell and bond data was exported. Using published matlab code^42^, we were able to infer tensions along individual cell edges. These tensions were then averaged across several cells to infer a map of stress (extracted from the eigenvectors of the inferred stress tensor) in the tissue. The eigenvectors define the orientation, and the eigenvalues define the magnitude of the lines.

#### ERK/Cdk quantification across tissue

Images acquired across multiple overlapping regions were stitched using Fiji. Custom code adapted from previous work^64^ was used to generate ERK/Cdk activity across the entire posterior trunk. Nuclei were segmented with TGMM using the equalized H2A-mCherry signal. A mask was generated using this nuclei channel and was used to measure nuclear signal intensity. This was dilated and subtracted to generate the cytoplasmic mask, which was used to measure the cytoplasmic signal intensity. The ratio of cytoplasm to nuclear signal was used to determine ERK/Cdk activity.

#### Single cell ERK/Cdk activity

Images were acquired sequentially for three channels and acquired every 30 mins. The movies were cropped and segmented using ilastik or trackmate plugin in Fiji. The tracking was verified manually and only those cells with proper tracks were used for further analysis. Mask was generated using nuclear channel and ERK/Cdk activity was determined by cytoplasmic to nuclear ratio.

#### Geminin proliferation counts

Images acquired from multiple overlapping regions were cleaned, flattened and stitched using Fiji. Auto local threshold was applied using Phansalkar method with radius 15 and cleaned to remove autofluorescence. Using custom matlab code, numbers of Geminin positive and total nuclei were quantified.

#### CWT analysis

Continuous Wavelet Transform (CWT) was performed using MATLAB’s cwt() function, with ERK activity traces spanning 20 datapoints or 5 hours being used as inputs for the cwt function. The resulting CWT heatmaps from individual ERK traces, which depict the magnitude of the pulsatile activity in the traces, were grouped based on 24-hour intervals, then averaged to produce one CWT heatmap. The cone of influence region was extracted from the CWT function and used to remove regions of the scalogram where the cwt analysis cannot be accurate. The color bar depicts the gradient of colors corresponding to the CWT magnitude.

## ACKNOWLEDGEMENTS

We would like to thank Akankshi Munjal and her lab for kindly sharing tools, reagents, and expertise. Authors would like to acknowledge Ashley Rich, Susanna Brantley, Priyom Adhyapok and Leslie Slota-Burtt for comments on the manuscript and members of Di Talia and Poss lab for their critical input and feedback during the project. NR was funded by NIH K01AR082432. SDT and KP acknowledge funding from NIH R01 AR076342.

## Author contributions

N.R., K.D.P. and S.D.T. conceived the project and designed the experiments; N.R. and M.O.B. conducted experiments and analyzed data; N.R., M.O.B., F.A.B, J.P., A.T.C., M.B. developed transgenic fish lines; C.R. developed computational tools and performed data analysis with help from N.R. and M.O.B.; N.R., K.D.P. and S.D.T., resources and funding acquisition; N.R., K.D.P. and S.D.T. wrote the paper. All authors read and commented on or edited the manuscript.

## Data and materials availability

All materials and codes are available upon request.

## Notes

### Competing Interest Statement

The authors have declared no competing interest.

