## Supplementary figure legend for "Phased ERK-responsiveness and developmental robustness regulate teleost skin morphogenesis"

**Supplementary Figure legends**

**Figure S1 – Neighbor exchange, cell size and shape are minor contributors to periderm expansion.**

(A) Stills from transgenic embryos expressing occludinb-GFP and H2A-mCherry under *krt4* promoter at 28 hpf. Scale bar – 100 µm.

(B) Quantification of number of cells undergoing neighbor exchange in zebrafish periderm cells and Drosophila germ band expansion, a process governed by convergent extension. Using our algorithm, we find <5% periderm cells undergo neighbor exchange, while about >20% cells undergo neighbor exchange in Drosophila tissue. N = 10 regions, 5 embryos – Zebrafish periderm.

(C) Example of Drosophila germ band cells tracked over time, showing a neighbor exchange event, marked by *, detected by our algorithm.

(D) Quantification of average cell area of zebrafish periderm cells covering the trunk region before and after live imaging. There is a moderate increase in cell area which could make a minor contribution towards periderm expansion. N = 5 embryos.

(E) Aspect ratio of zebrafish periderm cells covering the trunk region before and after live imaging, shows a trend towards decreasing during the imaging. N = 5 embryos.

(F) Histogram of tissue tension along A-P (Purple) vs D-V axis (blue), shows significantly increased stress along A-P.

(G) Quantification of orientation of long axis of periderm cells covering the trunk region. The angles are closer to 0, which suggests that cells are oriented along A-P axis both before (pink) and after imaging (blue), implying that cells do not reorient themselves for the rapid expansion. N= 5 embryos.

**Figure S2 – ERK-KTR sensor specifically reports MAPK mediated ERK activity in the periderm.**

(A) Stills from transgenic embryos expressing *krt4:*H2A-mCherry *and krt4:*Venus-hGeminin treated with DMSO or 2µM PD0325901 for 6-8 hrs. Embryos treated with PD03 have significantly decreased proliferation seen by reduction in geminin positive cells. Scale bar – 100µm.

(B) Quantification of fraction of geminin positive cells in embryos treated with DMSO or PD0325901, shows that PD03 treated embryos have significantly decreased proliferation as seen by reduction in normalized fraction of geminin positive cells. Average of DMSO = 1, n=12 embryos, Average PD03 = 0.6809, n=15 embryos. P<0.0001. This suggests that ERK activity is required for propagation through cell cycle.

(C) Maximum projection view of periderm cells covering the trunk region expressing *krt4:*ERK-KTR-mCerulean, treated with DMSO, 2µM PD0325901, 50 µM Roscovitine or both for 4-6 hours. The corresponding heatmaps of ERK activity shown in the bottom panel. Scale bar – 100µm.

(D) Quantification of average ERK activity in periderm cells covering the posterior trunk of embryos under different treatments. Average ERK activity significantly decreases upon PD0325901 treatment when compared to Roscovitine suggesting the KTR-sensor is mainly responsive ERK. Average ERK DMSO = 1.468, PD03 = 0.788, Roscovitine = 1.412 and both drug = 0.749.

(E) Logistic regression analysis to predict the probability of cell division based on the average ERK activity in 1-5hr period leading up to mitosis. AUC = 0.84

**Figure S3 – EGFR mediated ERK signaling regulates periderm growth**
(A) Schematic for working of Dronpa based optogenetic activation of ERK. Dronpa monomers are inserted at the N- and C-termini of constitutively active MEK-E203K. Activation with 405nm light induces Dronpa dimerization which sterically inhibits MEK activity. Photoconversion with 488nm light prevents Dronpa from dimerizing, leading to loss of its fluorescence and uncaging of active MEK which could then turn on ERK activity.
(B) Maximum projection view of periderm cells covering the trunk region expressing *krt4:*ERK-KTR-mCerulean, treated with DMSO, 10µM EGFR inhibitor - PD168393, 2 µM FGFR inhibitor - BGJ398 for 4-6 hours. The corresponding heatmaps of ERK activity shown in the bottom panel. Scale bar – 100 µm.
(C) Quantification of average ERK activity in periderm cells covering the posterior trunk of embryos under different treatment conditions mentioned in (B). Average ERK activity significantly decreases upon EGFR inhibitor treatment while only moderately decreases with FGFR inhibitor treatment, suggesting that ERK activity is the periderm covering posterior trunk is mediated by EGFR signaling. Average ERK activity upon DMSO treatment = 1.103, n= 7 embryos, average ERK activity upon EGFRi = 0.7883 and average ERK activity upon FGFRi = 0.9646. One-way ANOVA comparison DMSO vs EGFRi, $p<0.0001$ and DMSO vs FGFRi, $p = 0.0215$.
(D) and (E) Maximum intensity projection view of periderm cells expressing *krt4:*DronpaMEK203*, krt4:*H2A-mCherry and *krt4:*ERK-KTR-mCerulean*.* ERK activity visualized by ERK-KTR-mCerulean and corresponding heat maps below. In control regions (D) where Dronpa was not photoconverted, ERK activity remains comparable after 10 mins. Contrastingly, ERK activity dramatically increases in the same region, upon 10 mins of Dronpa photoconversion (E). Scale bar – 20µm.
(F) and (G) Quantification of fraction of geminin positive cells in embryos treated with DMSO or 10µM EGFR inhibitor - PD168393 for 6 and 18 hours respectively. EGFRi treated embryos have significantly decreased proliferation as seen by the reduction in normalized fraction of geminin positive cells at both time points. Average normalized fraction of geminin positive cells in DMSO at 6 hours post treatment (hpt) = 1, n= 6 embryos, Average EGFRi = 0.06, n= 6 embryos. $P<0.0001$, unpaired t-test. Average normalized fraction of geminin positive cells in DMSO at 18 hpt = 1, n= 7 embryos, Average EGFRi = 0.10, n= 8 embryos. $P=0.0007$.

**Figure S4 – Elongation rate of posterior trunk region of zebrafish embryos slows after 2 dpf.**
(A) Brightfield view of zebrafish embryos at 29, 46 and 74 hpf respectively. Dashed line indicates the anterior limit for our trunk length measurements. Scale bar – 500 µm.
(B) Quantification of rate of change in length of the posterior region (length from dashed line), shows that embryo elongation slows down to about half the rate after 58 hpf. Average rate of growth 28-46 hpf – $38.57 \pm 3.224$ µm/hr, n = 18 embryos and average rate of growth 52-70 hpf – $19.56 \pm 3.478$ µm/hr, n = 18 embryos. Growth rates are significantly different with $p< 0.0001$.

**Figure S5 – Cdk2 sensor specifically reports Cdk activity in the periderm.**

(A) Maximum projection view of periderm cells covering the trunk region expressing *krt4:*Cdk2-KTR-mCherry, treated with DMSO or 50 µM Roscovitine for 4-6 hours. High magnification of boxed regions, shows the shift in sensor localization from cytoplasmic to nuclear upon Cdk inhibition, suggesting that Cdk2 sensor is sensitive to Cdk activation levels. Scale bar – 100 µm.
(B) Quantification of percentage mitotic cells in periderm covering the posterior trunk of embryos under different treatment conditions mentioned in (A), shows reduction in mitotic cells upon Cdk inhibition. Percentage mitotic cells with DMSO = 32.60% and mitotic cells upon 50 µM Roscovitine = 21.90 %, N = 5 embryos each condition. p = 0.028 unpaired t test.
(C) Maximum projection view of periderm covering posterior trunk expressing *krt4:*Venus-hGeminin and *krt4:*H2A-mCherry at 28 and 52 hpf, showing the reduction in Geminin expressing cells at the latter time point.
(D) Quantification of percentage of cells expressing Geminin at 28-30 hpf and 52-54 hpf, shows significant reduction in Geminin expressing cells at 52-54 hpf. 28-30 hpf, n = 8 embryos, average = 35.65, 52-54 hpf, n = 8 embryos, average = 5.655. Welch t-test, p < 0.0001.

**Figure S6 – Shift in proliferative to non-proliferative response to ERK signaling as periderm expansion slows**
Scatter plot of ERK and Cdk activities in single cells of the periderm at 30-50 hpf (A). The percentage of cells with high ERK activity and high Cdk activity (blue population) is 21 % of total cells, while that of high ERK activity and low Cdk activity (pink population) is 14% of total cells. High ERK is denoted by cells with ERK activity >=1. N = 4 embryos.

**Figure S7 – Periderm growth exhibits adaptive robustness**
(A) Brightfield view of control and p21 expressing zebrafish embryos at 28 hpf, 48 hpf and 72 hpf dpf respectively. Dashed line indicates the anterior limit for our trunk length measurements. Scale bar – 500 µm.
(B) Quantification of rate of change in length of the posterior region (length from dashed line), shows that embryo elongation occurs in *krt4:*p21-2a-GFP transgenic embryo. Further, it slows down to about half the rate after 48 hpf, similar to control embryos. Average rate of growth 28 to 46 hpf control embryo– 38.57 µm/hr, n = 18 embryos and *krt4:*p21-2a-GFP embryos – 41.52 µm/hr, n = 17 embryo, p = 0.038. Average rate of growth 52 to 70 hpf control embryos – 19.56 µm/hr, n = 18 embryos and *krt4:*p21-2a-GFP embryos – 17.33 µm/hr, n = 17 embryos, p = 0.33. multiple comparison, One-way ANOVA.
(C) Quantification of coefficient of variation of cell size in WT control and p21 expressing larvae at 48 hpf, shows higher variability in p21 expressing periderm cells. Average coefficient of variation in WT – 0.36, n = 9, p21 expressing larvae – 0.43, n = 8, p = 0.0381, unpaired t-test.
(D) Quantification of coefficient of variation of cell size in WT control and p21 expressing adults at 6 months, shows variability in cell size is comparable between p21 expressing embryos and WT embryos. Average coefficient of variation in WT – 0.27, n = 6, p21 expressing larvae – 0.31, n = 6; P = 0.1791, unpaired t-test.
