## Supplementary figures for "Phased ERK-responsiveness and developmental robustness regulate teleost skin morphogenesis"

Fig S1 - Neighbor exchange, cell size and shape are minor contributors for periderm expansion

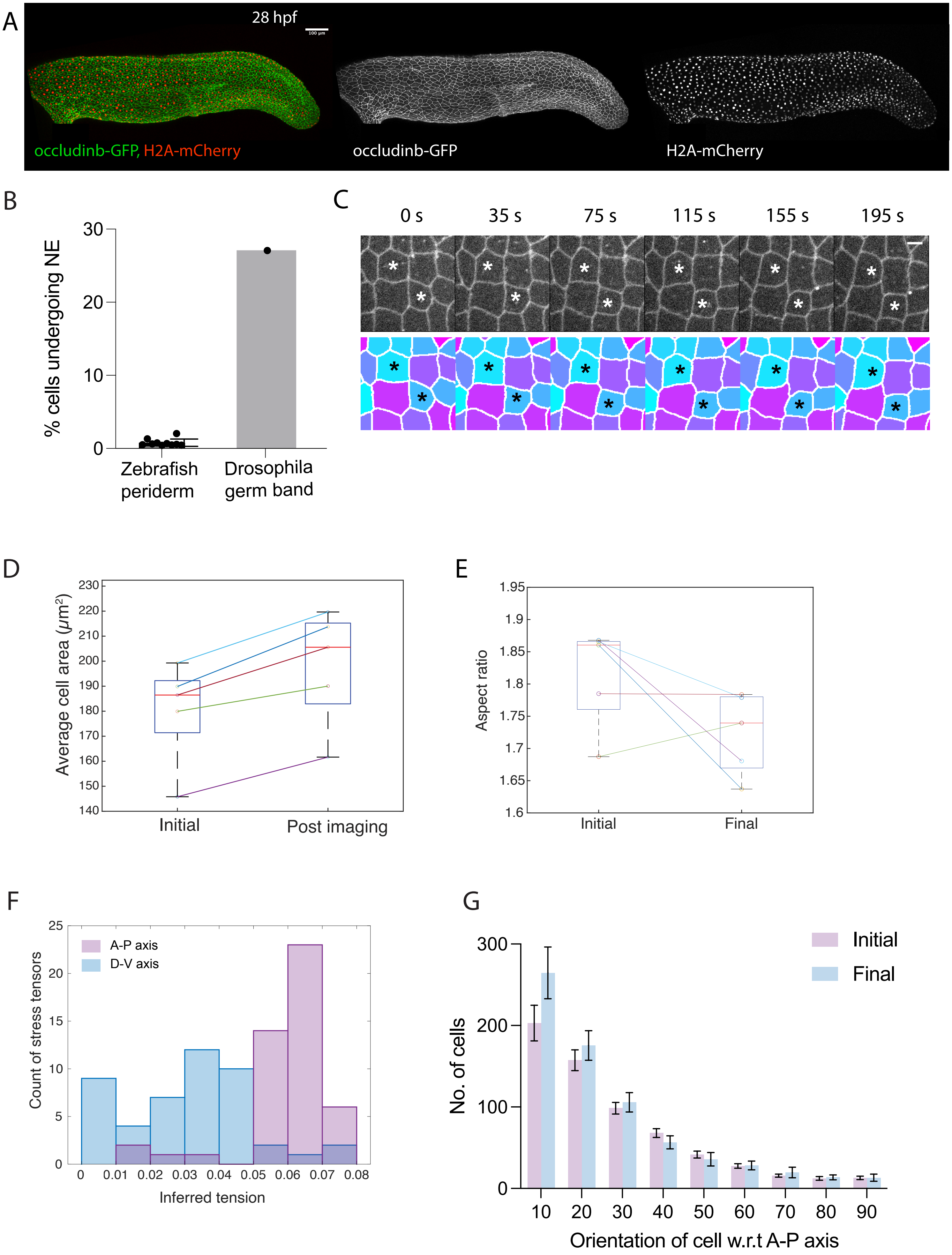

Fig S2 - ERK-KTR sensor specifically reports MAPK mediated ERK activity in the periderm

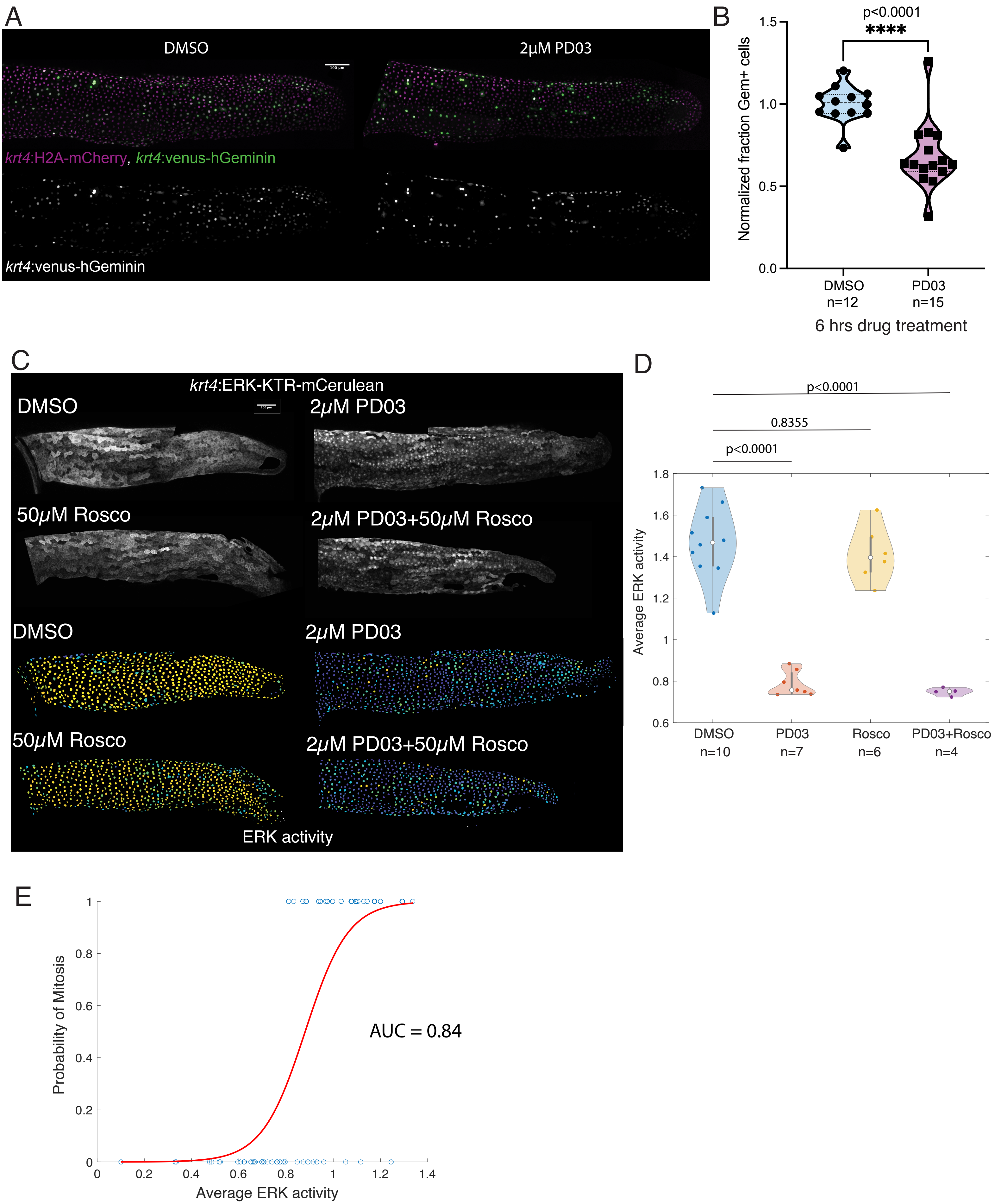

Fig S3 - EGFR mediated ERK signaling regulates periderm growth

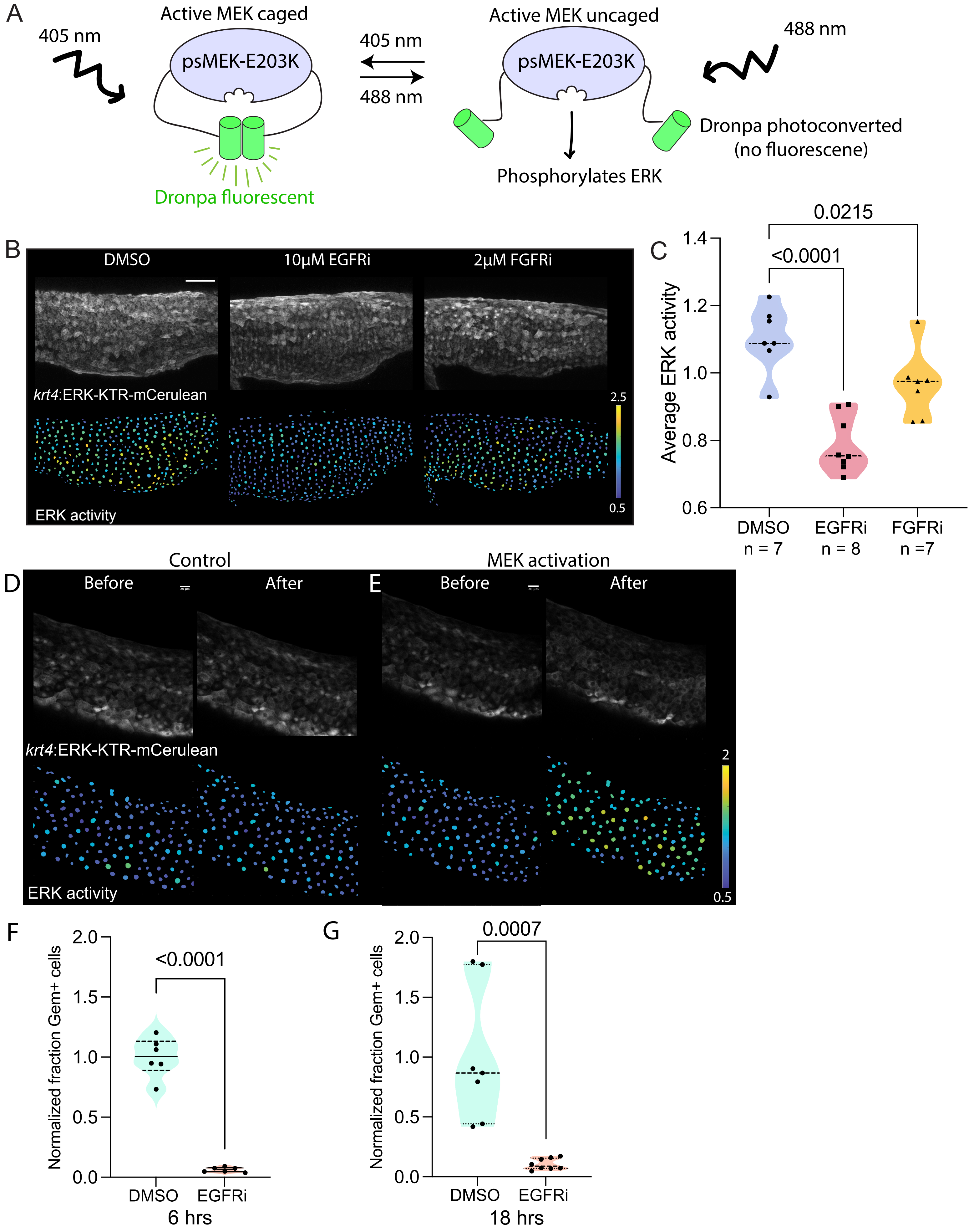

Fig S4 - Elongation rate of posterior trunk region of zebrafish embryo slows down after 2 dpf

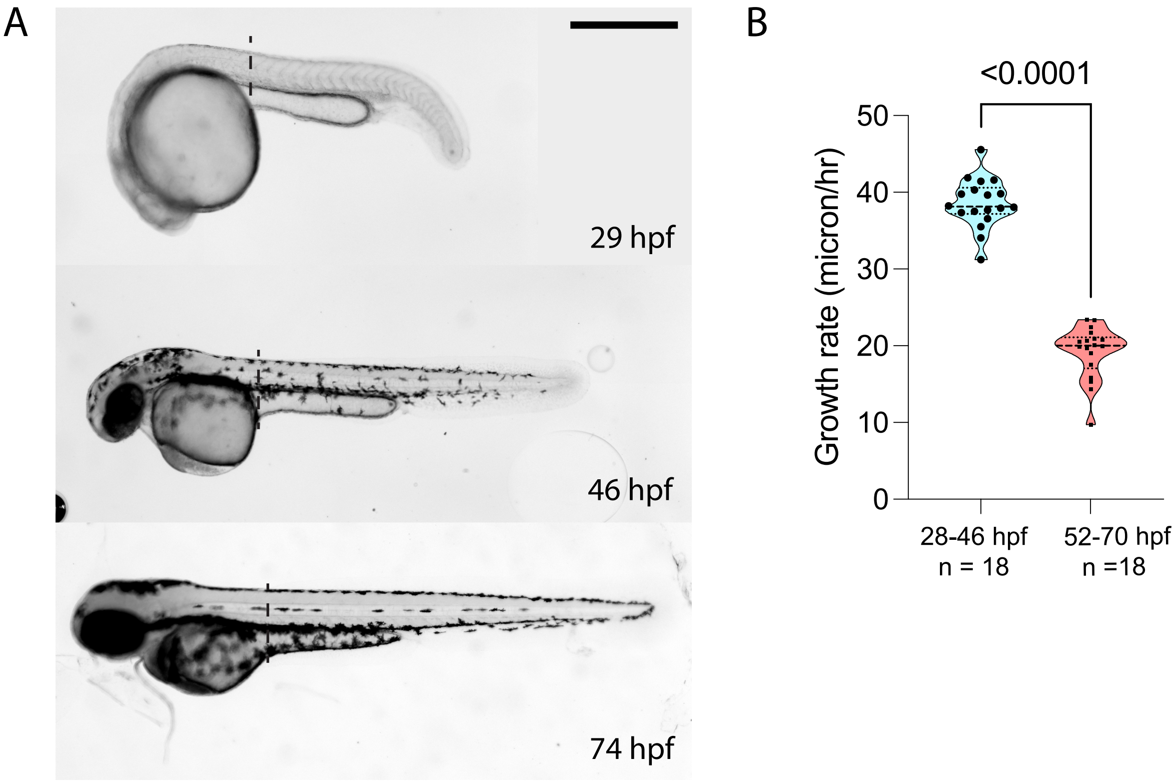

Fig S5 - Cdk2 sensor specifically reports Cdk activity in the periderm

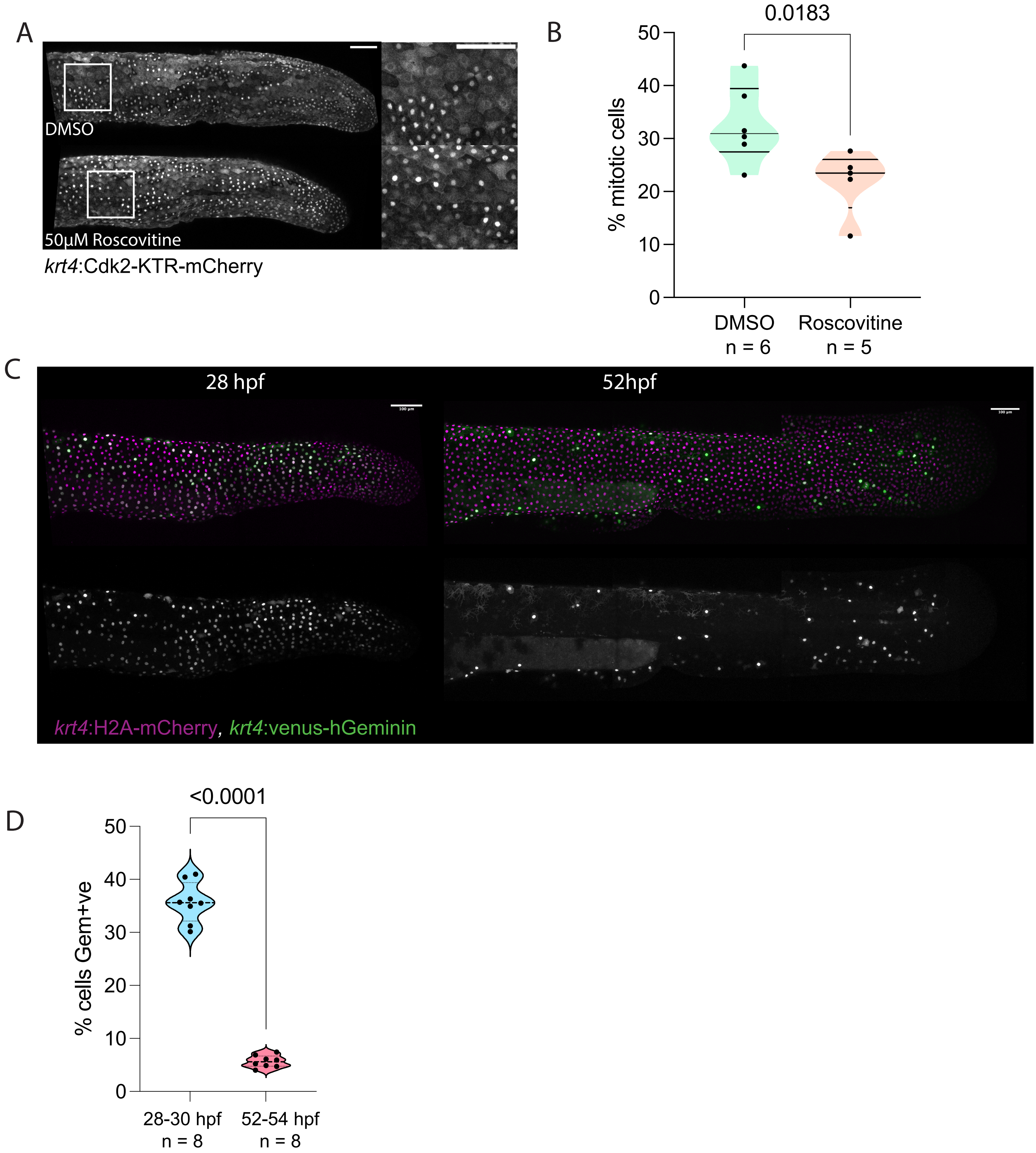

Fig S6- Shift in proliferative to non-proliferative response to ERK activity as elongation rate dampens

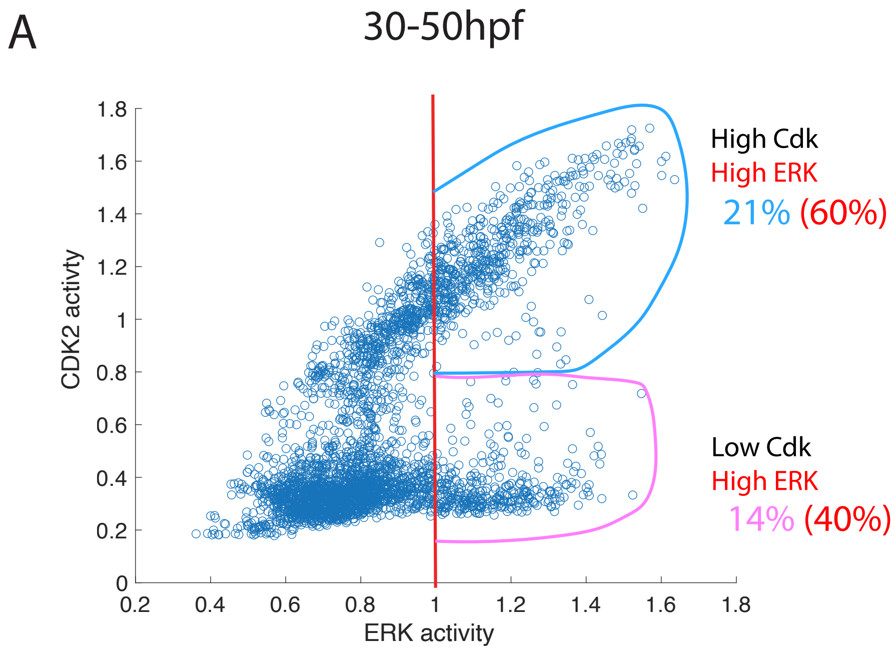

Figure S7 - Axial elongation occurs normally when periderm proliferation is inhibited

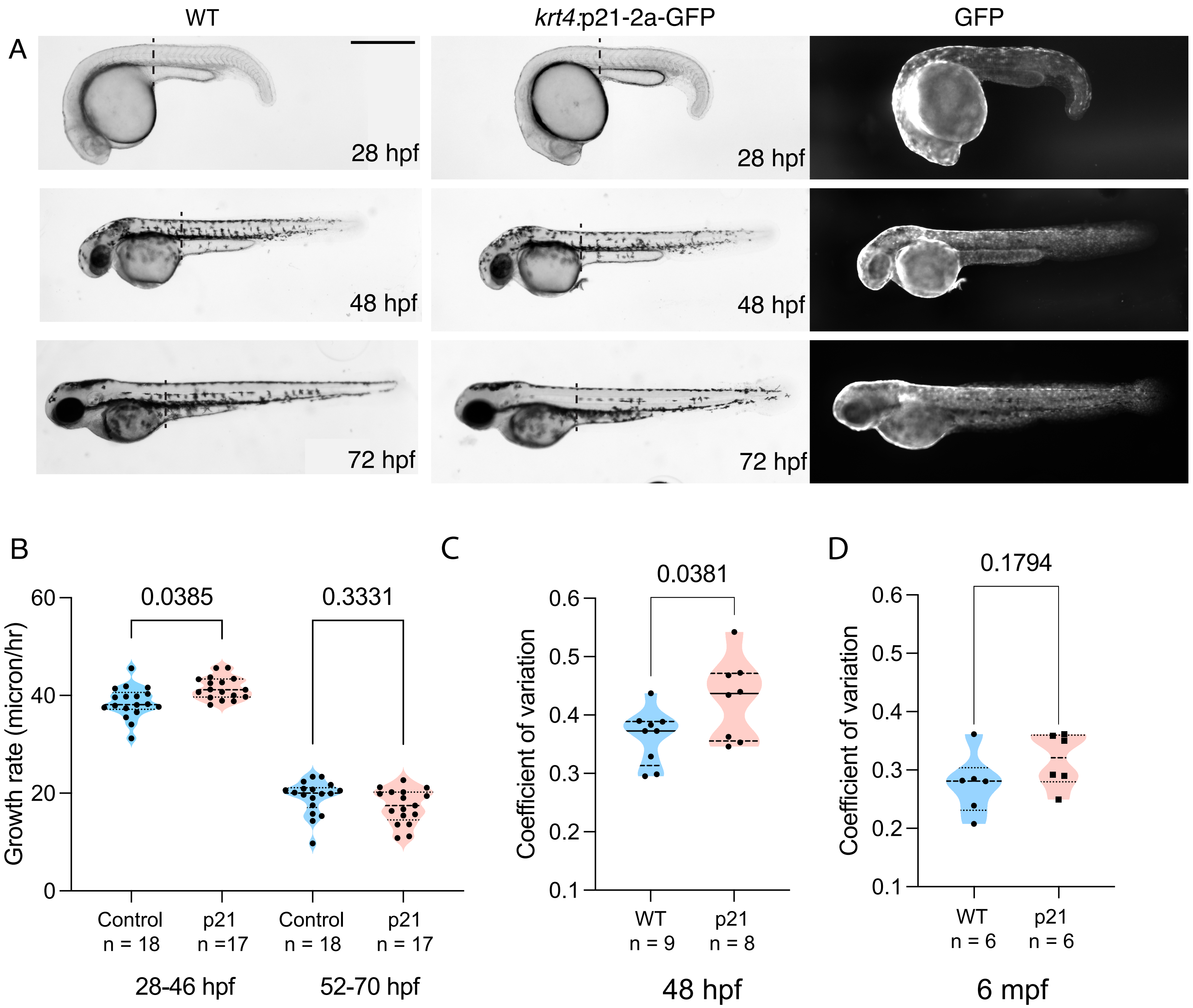
